# Sex Differences in DNA Methylation and Gene Expression Reveal Distinct Molecular Programs in Muscle-Invasive Bladder Cancer

**DOI:** 10.64898/2026.10.21.760777

**Authors:** Bhavisha Doshi, Eduardo Cortes Gomez, Barbara A. Foster, Jianmin Wang, Norbert Sule, Wendy J. Huss, Anna Woloszynska

**Author notes:** Corresponding author; email: Department of Pharmacology and Therapeutics, Roswell Park Comprehensive Cancer Center, Buffalo, NY, 14263, United States of America. **Acronyms** Muscle-invasive bladder cancer (MIBC), N-butyl-N-(4-hydroxybutyl) nitrosamine (BBN), epithelial-mesenchymal transition (EMT), cytosine-phosphate-guanine dinucleotide (CpG), X-chromosome inactivation (XCI), tumors (T), tumor-adjacent normal tissues (NAs/NT), Gene Expression Omnibus (GEO), Gene set enrichment analysis (GSEA), differentially methylated regions (DMRs), differentially expressed genes (DEGs), Maxdiff (from the methylation array data), log2Fold Change (log2FC), transitional cell carcinoma (TCC), invasive urothelial carcinomas (iUC), including tumors with squamous differentiation (iUCSD) and sarcomatoid features (iUCsarc).

## Abstract

Sex differences in muscle-invasive bladder cancer (MIBC) incidence and outcomes are well established, but the underlying epigenetic mechanisms remain poorly understood. We integrated genome-wide DNA methylation and RNA sequencing to define sex-associated epigenetic and transcriptional programs in human MIBC and evaluated their conservation in the N-butyl-N-(4-hydroxybutyl) nitrosamine (BBN) mouse model. Male tumors exhibited substantially broader DNA methylation remodeling, whereas female tumors showed prominent epithelial-mesenchymal transition (EMT) and mitotic programs. These sex-associated patterns were recapitulated in BBN tumors, with substantial conservation of human MIBC methylation changes and transcriptional programs. Cross-species integration identified conserved methylation-regulated genes, including *ADAMTS12*, *ONECUT2*, and *PPP2R2C*, whose expression increased following pharmacologic DNA demethylation and was associated with disease progression. Given the potential role of sex chromosomes in sex-associated molecular differences in MIBC, we evaluated X-chromosome inactivation escape and Y-linked gene expression. Reduced expression of an X-chromosome inactivation escape signature was independently associated with shorter overall survival, while reduced Y-linked gene expression was associated with immune activation. Together, these findings identify conserved sex-associated epigenetic programs in MIBC and demonstrate distinct methylation, transcriptional, and sex-chromosome features that may contribute to sex differences in bladder cancer biology and clinical outcomes.

## Background

Sex-based differences in muscle-invasive bladder cancer (MIBC) are well established. Men have an approximately four times higher incidence, while female patients experience poorer outcomes^1,2^. Women are also more likely to be diagnosed at a later stage, which may contribute to their less favorable disease course^3^. Beyond differences in stage at diagnosis, men and women also exhibit distinct patterns of metastatic spread^4^. Men show a greater age-related increase in bone and brain metastases, whereas women experience a higher age-related increase in lung and liver metastases^4^. These differences underscore that sex influences multiple aspects of tumor behavior in bladder cancer.

Bladder cancer is a major global health concern that ranks as the ninth most common cancer worldwide^5^ with MIBC accounting for up to 30% of new diagnoses^6^. MIBC has a 5-year survival rate of 39% for lymph node-positive tumors, which decreases to only 8.8% for distant metastases^7^. Several risk factors contribute to the development of bladder cancer, including smoking, age, sex, and race^1^. Smoking is the primary risk factor, significantly increasing the likelihood of developing the disease^8^. Tobacco exposure is estimated to account for approximately 50-65% of bladder cancer cases in men and 20-30% in women^8^. Although the higher incidence in men has long been attributed to higher smoking rates^9^, the risk remains elevated in men even after adjusting for smoking status^8^. The persistence of these sex differences suggests that biological factors, in addition to environmental exposures, contribute to disease variability. Our previous work has summarized potential mechanisms underlying these disparities, highlighting the complexity of sex-associated tumor biology^1^.

Bladder cancer arises from a complex interplay of genetic and epigenetic alterations that drive tumor initiation and progression^10–12^. Although somatic mutations in cancer susceptibility genes contribute to tumorigenesis, mounting evidence highlights DNA methylation as a pivotal epigenetic mechanism disrupted in bladder cancer^13,14^. DNA is methylated by the covalent addition of a methyl group to the 5-position carbon of cytosine within a cytosine-phosphate-guanine (CpG) dinucleotide^15^. Promoter CpG island methylation is generally associated with transcriptional repression, whereas methylation within gene bodies and distal regulatory regions, including CpG shores, shelves, and open-sea regions, can have context-dependent effects on transcription and chromatin regulation^16^. The bladder cancer methylome is characterized by global hypomethylation coupled with focal hypermethylation in CpG-rich regions, leading to dysregulated expression of tumor suppressor genes and oncogenes^10,17^. Targeting DNA methylation regulators has shown promise in enhancing cisplatin-based neoadjuvant chemotherapy, the first-line treatment for MIBC, by improving therapeutic response and re-sensitizing cisplatin-resistant tumors^18–20^. Together, these findings highlight the relevance of DNA methylation to both bladder cancer biology and therapeutic response. Previous studies examining sex-associated DNA methylation in bladder cancer have been limited and focused on individual targets, including *LINE1* and *TOP2B* ^21,22^. More recently, a pan-cancer analysis of TCGA data identified sex-associated differences in DNA methylation remodeling in bladder cancer, including a greater magnitude of methylation changes in males^23^. However, the relationship between sex-associated methylation remodeling and transcriptional programs in MIBC remains largely unexplored.

The N-butyl-N-(4-hydroxybutyl)-nitrosamine (BBN) model is a widely used and robust mouse model for studying bladder cancer^24,25^. BBN is a bladder carcinogen that models aspects of exposure to N-nitroso compounds present in tobacco smoke^26^. Following BBN exposure, mice develop invasive bladder tumors that exhibit histologic and molecular characteristics similar to human MIBC, including high mutation burden and expression of basal cancer markers^24–26^. However, it remains unclear whether BBN-induced bladder tumors recapitulate DNA methylation changes observed in human bladder cancer.

The objective of our study was to define sex-associated differences in DNA methylation and gene expression in MIBC and to identify the biological pathways underlying these differences. We integrated genome-wide DNA methylation and RNA sequencing data to identify DNA methylation-associated transcriptional changes and biological pathways in MIBC. We further examined whether these epigenetic and transcriptional features were conserved in BBN-induced bladder tumors. We also examined X-chromosome inactivation escape and Y-chromosome gene expression as potential contributors to sex-associated tumor biology. Our analyses revealed distinct sex-associated DNA methylation and transcriptional programs, with broader methylation remodeling in male tumors and distinct EMT-associated pathway enriched in female tumors compared to their male counterparts. Together, our findings identify distinct and conserved epigenetic and transcriptional programs in male and female bladder cancer that may contribute to sex differences in MIBC biology and clinical outcomes.

## Results

### Tumor status and sex contribute to molecular variation in bladder cancer

To define the major sources of molecular variation in bladder cancer, we first characterized DNA methylation in the Roswell Park bladder cancer cohort^27^, which included primary muscle-invasive bladder tumors (T) and tumor-adjacent normal tissues (NAs/NT) from both male and female patients. Principal component analysis (PCA) of DNA methylation across all samples showed clear separation between tumor and NAs/NT (Figure 1A). Eigenvector correlation analysis identified tumor status (i.e. T vs Nas/NT) as strongly associated with PC1 and PC2, indicating that tumor status was the predominant source of methylation variation across the cohort (Figure 1B).

**Figure 01.**
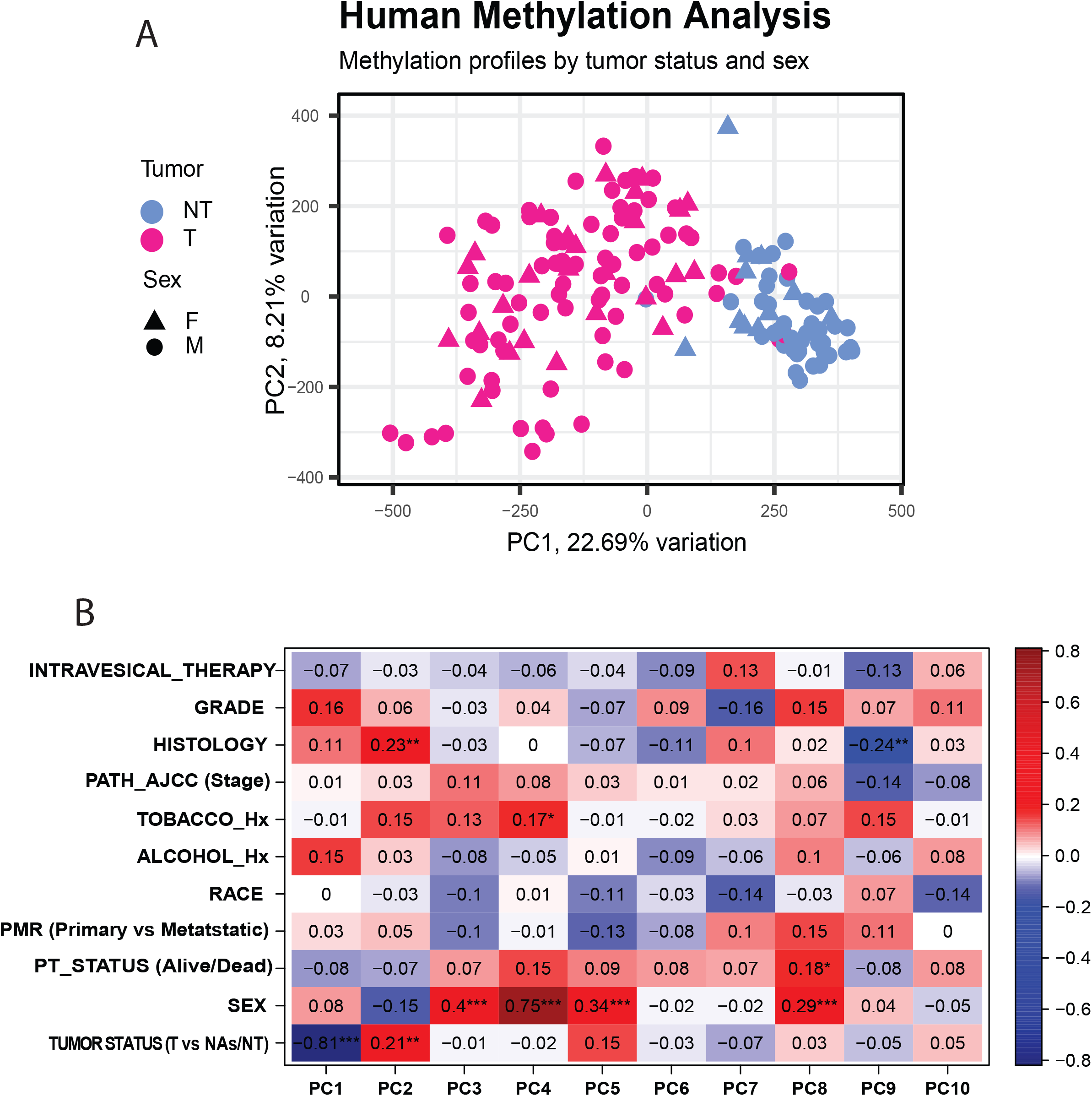
DNA methylation profiles distinguish tumor status and sex in MIBC. (A) Principal component analysis (PCA) of DNA methylation profiles from MIBC tumors (n = 105) and tumor- adjacent normal tissues (n = 52) in the Roswell Park cohort. Samples are plotted according to the first two principal components, with each point representing an individual sample. (B) Eigenvector correlation analysis identifying clinical and biological variables associated with principal components PC1-PC10. Each cell represents the correlation between the indicated clinical variable and the corresponding principal component. The color scale represents the direction and magnitude of the correlation, with values closer to +1 indicating a strong positive correlation, values closer to -1 indicating a strong negative correlation, and values near 0 indicating little or no correlation. Asterisks indicate statistically significant correlations, with P < 0.05, P < 0.01, P < 0.001, and P < 0.0001. (Acronyms: Path_AJCC -Pathological American Joint Committee on Cancer staging, Tobacco_HX - Tobacco history, Alcohol_HX – Alcohol history, PMR - Primary or metastatic disease, PT_Status - Patient status).

Tumor histology was also associated with PC2, prompting us to further stratify the samples by histological subtype (Figure 1B). Notably, patient sex was associated with PC3, identifying sex as an additional source of variation in the bladder tumor methylome (Figure 1B). We next examined whether associations of tumor status, sex and histology were also evident at the transcriptional level. RNA-seq analysis similarly showed clear separation between tumors and NAs/NT (Supplementary Figure 1A), with tumor versus adjacent normal tissue ROS131-34 (T vs NT/Nas) strongly associated with PC1, PC2 and PC3 (Supplementary Figure 1B). Sex was associated with PC13 (Supplementary Figure 1B). Together, these analyses identify tumor status as a dominant source of molecular variation across both DNA methylation and gene expression, with sex contributing an additional source of molecular variation.

### Sex-specific methylation and gene expression profiles reveal distinct molecular programs in bladder cancer

To characterize sex-associated molecular differences in MIBC, we analyzed DNA methylation and RNA-seq from the Roswell Park bladder cancer cohort^27^. We focused on transitional cell carcinoma (TCC), the predominant histological subtype of MIBC, which accounts for approximately 90% of bladder cancer cases^28^. The DNA methylation cohort comprised 44 male tumors and 26 matched NAs, and 18 female tumors and 7 matched NAs. RNA-seq analysis included 29 male TCC tumors with 7 matched NAs and 10 female TCC tumors with 3 matched NAs.

Sex-stratified DNA methylation analysis revealed substantially more extensive DNA methylation changes in male compared to female tumors relative to their respective NAs. Male tumors exhibited 11,097 hypomethylated and 11,498 hypermethylated genes, compared with 852 hypomethylated and 906 hypermethylated genes in female tumors (Supplementary Tables 2 and 3).

We next examined transcriptional changes within each sex. Relative to NAs, male tumors exhibited 3,850 upregulated and 2,484 downregulated genes, whereas female tumors exhibited 957 upregulated and 1,198 downregulated genes (Supplementary Figures 2A and 2B). Gene set enrichment analysis (GSEA) identified distinct pathway-level changes in male and female tumors. Female tumors showed enrichment of epithelial-to-mesenchymal transition (EMT) and mitotic spindle programs and reduced enrichment of the UV response, whereas male tumors showed enrichment of inflammatory response, interferon alpha response, and androgen signaling (Supplementary Figure 2C).

**Figure 02.**
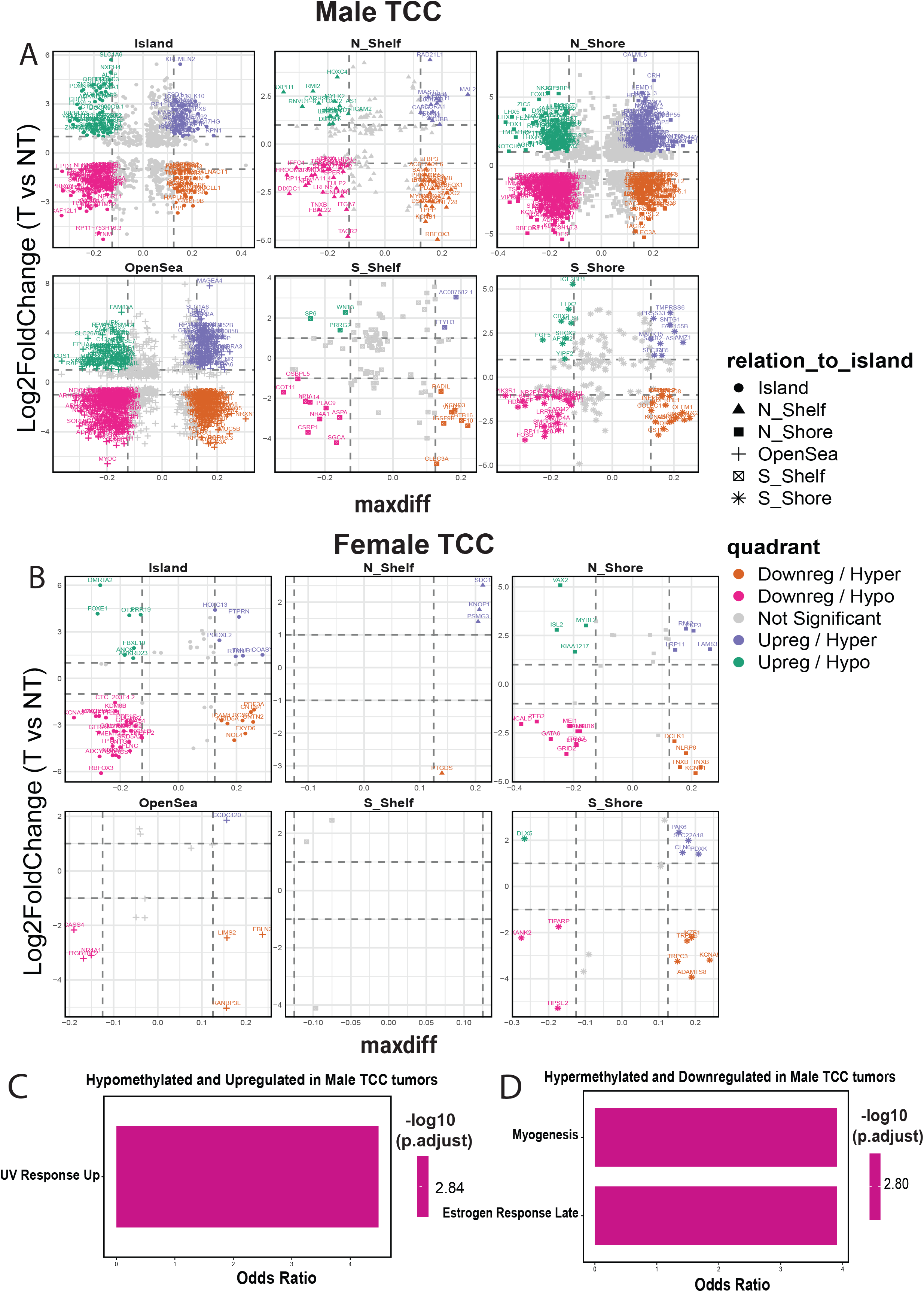
Human TCC exhibits sex-specific epigenetic and transcriptional landscapes. (A) Faceted scatter plots showing methylation regions (CpG island, N_shelf, N_shore, OpenSea, S_shelf, S_shore) with integrated significant DMRs (p < 0.05, maxdiff threshold) and significant DEGs (p < 0.05, log2 fold-change threshold) in male TCC. Quadrants indicate combined methylation-expression patterns: upregulated/hypermethylated (top-right), upregulated/hypomethylated (top-left), downregulated/hypermethylated (bottom-right), and downregulated/hypomethylated (bottom-left). (B) Faceted integration of significant DMRs and DEGs in female TCC, with quadrant definitions as described in panel A. (C) Pathway enrichment of hypermethylated and downregulated genes in male TCC using ENRICHR hallmark gene sets. Bar colors indicate -log10(p-values). (D) Pathway enrichment of hypomethylated and upregulated genes in male TCC. Bar colors indicate -log10(p-values).

To identify transcriptional changes associated with differential DNA methylation, we integrated differentially methylated regions (DMRs) with differentially expressed genes (DEGs). Sex-stratified analysis identified distinct methylation-associated gene sets in male and female tumors. Among the methylation-associated genes identified in male tumors, 277 were hypermethylated and downregulated and 227 were hypomethylated and upregulated, with associated DMRs distributed across CpG islands, shores, shelves, and open-sea regions (Figure 2A). In comparison, 22 genes in female tumors were hypermethylated and downregulated and 12 were hypomethylated and upregulated (Figure 2B). Pathway analysis of these methylation-associated genes showed enrichment of UV response up pathway among hypomethylated and upregulated genes in male tumors (Figure 2C), whereas hypermethylated and downregulated genes in male tumors were enriched for myogenesis and estrogen response late pathways (Figure 2D). In the female cohort, the number of methylation-associated genes was insufficient to identify significantly enriched pathways.

Together, these analyses reveal sex-associated differences in the molecular changes accompanying MIBC, including substantially more extensive DNA methylation alterations in male tumors and distinct transcriptional programs in male and female tumors.

### XCI gene expression is associated with tumor aggressiveness and clinical outcome

Sex chromosomes represent a potential source of sex-associated molecular variation in bladder cancer. XCI escape genes are expressed from both X chromosomes in females but from a single X chromosome in males, resulting in sex differences in gene dosage ^29,30^. We therefore examined whether expression of genes that escape X chromosome inactivation (XCI) or genes encoded on the Y chromosome was associated with clinical outcome and tumor biology. We generated an XCI escape gene expression signature and evaluated its association with overall survival in a model adjusted for sex and tumor stage. Patients with low XCI escape signature expression had significantly shorter overall survival than those with high expression (p = 0.04; Figures 3A and 3C). When analyzed separately by sex, the association was not significant in females (p = 0.66) or males (p = 0.064) (Supplementary Figures 3A-D), potentially reflecting the greater range of XCI escape gene expression in the combined cohort.

**Figure 03.**
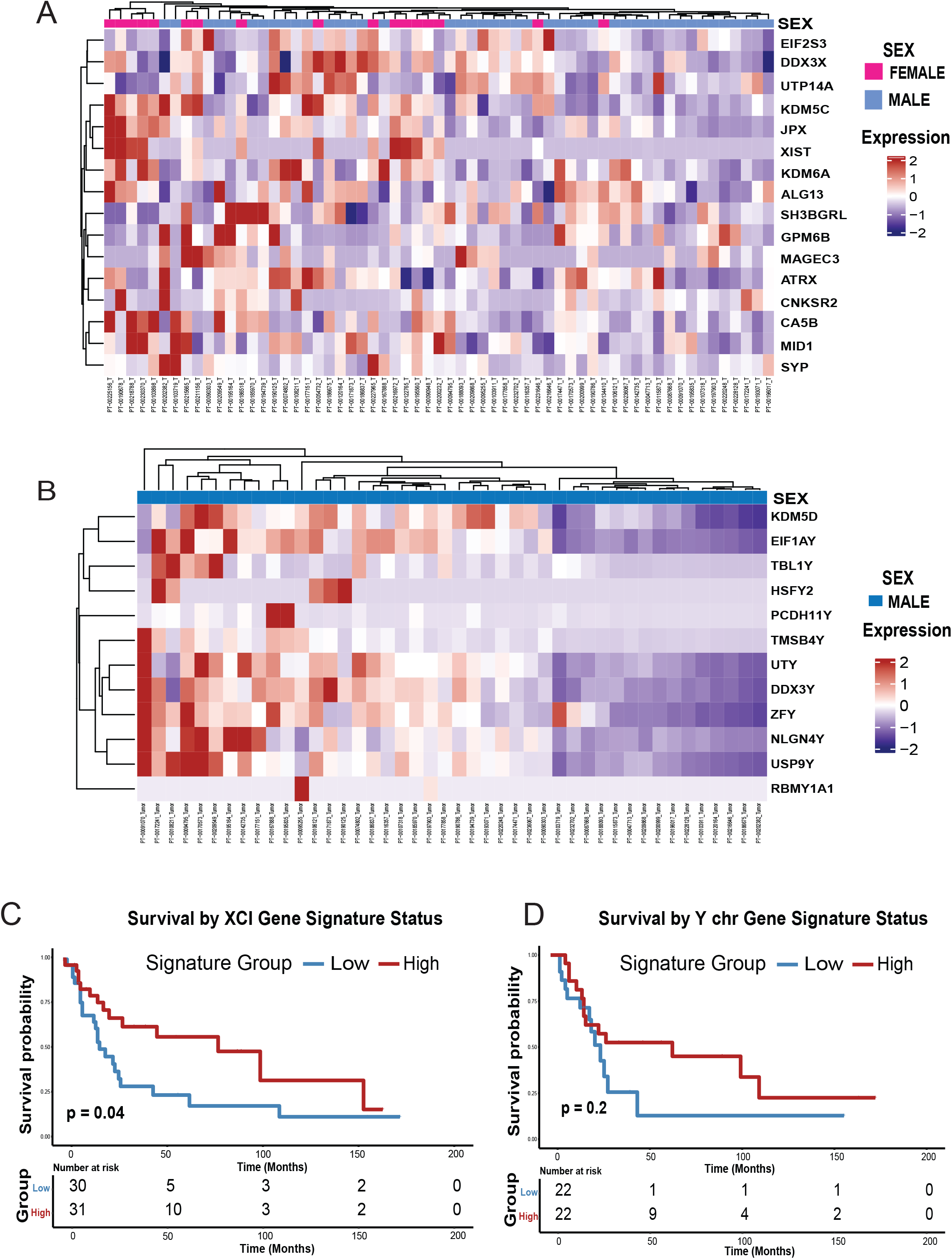
Expression of X chromosome escape genes is associated with clinical outcomes in bladder cancer. (A) Unsupervised clustering of genes that escape X-chromosome inactivation (XCI) across human TCC samples. Expression is shown using a red-to-blue gradient, with red representing upregulation and blue representing downregulation. Sample sex is annotated, with pink for female and blue for male samples. (B) Unsupervised clustering of Y chromosome genes across human male TCC samples. Expression is shown using a red-to-blue gradient, as in panel A. (C) Kaplan-Meier survival analysis comparing patients with low versus high XCI signature expression, generated using the Survminer package in R. Low-expression patients are shown in blue, and high-expression patients in red. (D) Kaplan Meier survival analysis comparing patients with low versus high Y chromosome gene signature expression. Low-expression patients are shown in blue, and high-expression patients in red.

We next examined Y chromosome gene expression in male tumors. The Y chromosome gene expression signature showed substantial interpatient variation, including a subset of tumors with markedly reduced Y-linked gene expression (Figure 3B), but was not significantly associated with overall survival (Figure 3D).

To determine whether sex chromosome gene expression was associated with distinct tumor transcriptional states, we compared tumors with high versus low XCI escape and Y chromosome signatures. Tumors with low XCI escape expression showed increased expression of several genes associated with tumor progression and metastasis, including *KRT6B*, *KRT6C*, *SPINK4*, and *ROS1*^31–34^ (Figure 4A). Similarly, tumors with low Y chromosome expression showed increased expression of tumor-progression-associated genes, including *CALML5* and *KRT6B*^34,35^ (Figure 4B).

**Figure 04.**
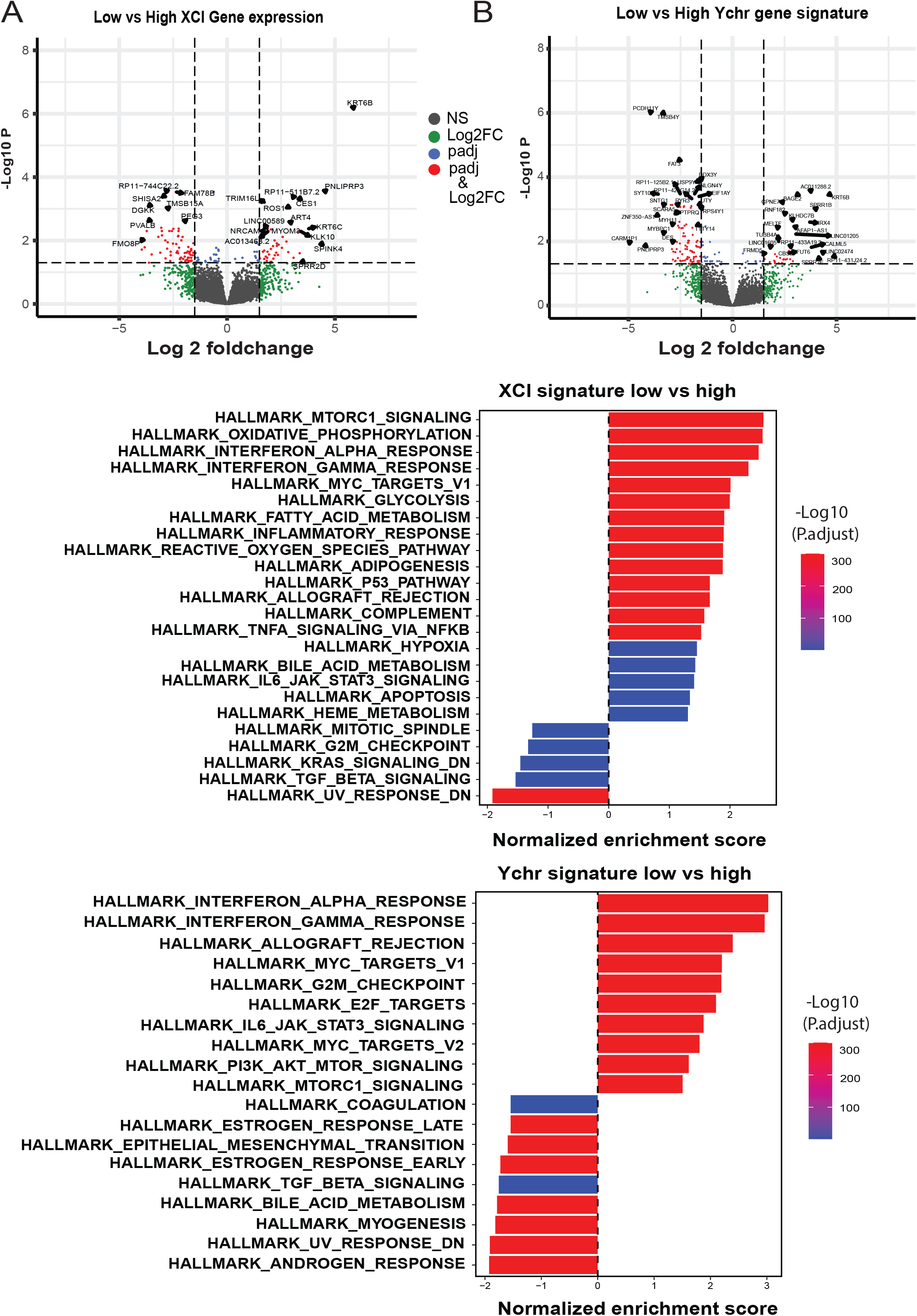
Low XCI escape and Y chromosome gene expression associate with aggressive and immune related tumor programs. (A) Volcano plot of differentially expressed genes (DEGs) identified by DESeq2 between patients with low versus high XCI signature expression. Genes with adjusted p < 0.05 are highlighted in red. (B) Volcano plot of DEGs identified by DESeq2 between patients with low versus high Y chromosome signature expression. Genes with adjusted p < 0.05 are highlighted in red. (C) Pathway enrichment analysis of DEGs associated with low versus high XCI signature expression using hallmark gene sets. Bar color represents adjusted p-value, and the X-axis shows normalized enrichment score (NES). (D) Pathway enrichment analysis of DEGs associated with low versus high Y chromosome signature expression, shown as in panel C.

GSEA further identified distinct biological programs associated with XCI and Y chromosome expression. Tumors with low XCI escape signature were enriched for MTORC1 signaling, oxidative phosphorylation, MYC target, and reactive oxygen species pathways (Figure 4C), consistent with oncogenic and metabolic programs associated with tumor growth and survival. Tumors with low Y chromosome expression were enriched for interferon alpha and interferon gamma response pathways, whereas tumors with high Y chromosome expression showed enrichment of epithelial-to-mesenchymal transition (EMT) and TGF-β signaling (Figure 4D). These findings are consistent with previous reports linking Y chromosome loss in bladder cancer with enhanced response to anti-PD-1 immune checkpoint blockade^36^ and further suggest that Y chromosome expression is associated with distinct immune and tumor-state programs.

Together, these findings identify sex chromosome gene expression as a feature of clinically and biologically distinct MIBC subsets. Low XCI escape gene expression was associated with poorer overall survival and enrichment of oncogenic and metabolic programs, whereas Y chromosome expression was associated with distinct immune and EMT-related transcriptional states but not overall survival.

### Carcinogen-induced mouse bladder tumors recapitulate molecular features of human MIBC

To determine whether molecular features identified in human MIBC are conserved in an experimentally tractable model, we examined the N-butyl-N-(4-hydroxybutyl) nitrosamine (BBN)-induced mouse model. BBN tumors recapitulate key histopathological and genomic features of human MIBC, but conservation of the tumor methylome has not been established. Male and female C57BL/6 mice were administered 0.1% BBN in drinking water and developed invasive urothelial carcinomas (iUC), including tumors with squamous differentiation (iUCSD) and sarcomatoid features (iUCsarc) (Supplementary Table 4). Because urothelial carcinoma with squamous differentiation is frequently associated with advanced disease, with approximately 60-70% of cases presenting as muscle-invasive bladder cancer, iUCSD tumors were selected for cross-species molecular analyses to minimize histological heterogeneity and facilitate comparison with human MIBC^37^.

We first compared DNA methylation profiles between human TCC tumors and NAs/NT with BBN-induced mouse tumors and normal mouse bladder. We identified 7,127 DMRs shared between human TCC and BBN-induced tumors, including 2,226 hypomethylated and 1,766 hypermethylated genes (Supplementary Figure 4A; Supplementary Table 5). Conserved hypomethylated genes were enriched for myogenesis, apical junction, epithelial-to-mesenchymal transition (EMT), and UV response down (Figures 5A and 5B), demonstrating conservation of tumor-associated epigenetic programs across species.

**Figure 05.**
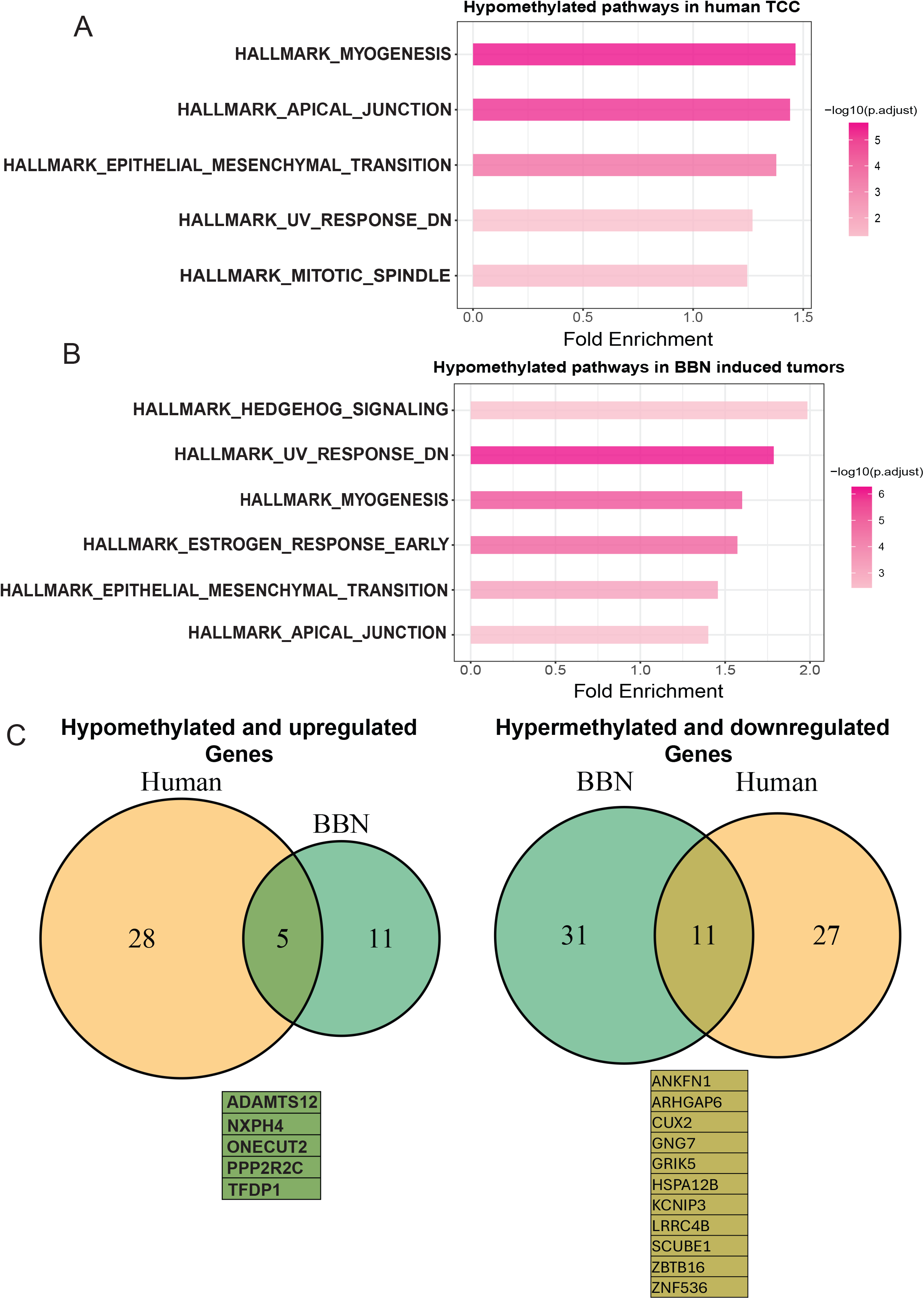
BBN-induced bladder tumors recapitulate the epigenetic and transcriptional landscapes of human MIBC. (A) Pathway enrichment analysis of hypomethylated genes in human TCC, performed using overrepresentation analysis with hallmark gene sets in R. The gradient of pink represents -log10 (adjusted P-values), and the X-axis indicates fold enrichment. (B) Pathway enrichment analysis of hypomethylated genes in BBN induced bladder tumors, performed using overrepresentation analysis with hallmark gene sets in R. (C) Venn diagram showing the overlap between human TCC and BBN-induced tumors for hypomethylated/upregulated genes (left) and hypermethylated/downregulated genes (right).

Integration of DMRs with DEGs identified five genes, *ADAMTS12*, *NXPH4*, *ONECUT2*, *PPP2R2C*, and *TFDP1,* that were consistently hypomethylated and upregulated in both human TCC and BBN-induced tumors, along with 11 genes that were hypermethylated and downregulated across species (Figure 5C; Supplementary Figures 5A and 5B). These findings demonstrate conservation of both tumor-associated DNA methylation changes and associated transcriptional alterations.

We next examined conservation of sex-associated molecular features. DNA methylation profiling included 8 male and 6 female iUCSD tumors and normal bladder tissues from 2 male and 2 female mice. Male BBN tumors exhibited 4,052 hypermethylated and 3,616 hypomethylated genes, whereas female tumors exhibited 939 hypermethylated and 436 hypomethylated genes (Supplementary Tables 6 and 7). Consistent with the broader methylation remodeling observed in human male tumors, 593 hypomethylated and 574 hypermethylated genes were shared between human and BBN male tumors. In contrast, only 11 hypermethylated and 1 hypomethylated genes were shared between human and BBN female tumors (Figures 6A and 6B).

**Figure 06.**
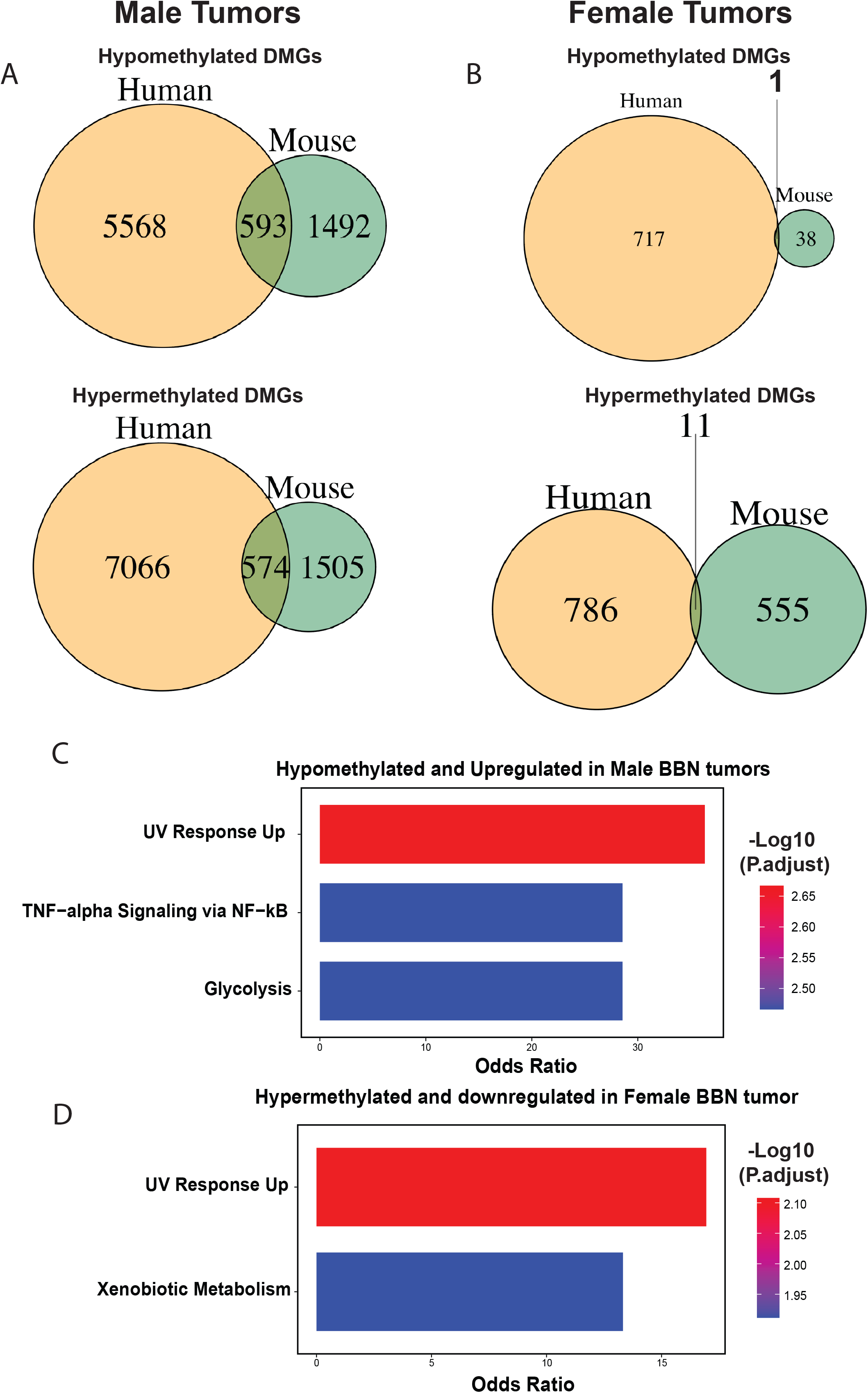
BBN-induced bladder tumors recapitulate the sex-specific epigenetic and transcriptional landscapes of human MIBC. (A) Venn diagram showing the overlap of hypomethylated genes (top) and hypermethylated genes (bottom) between male human TCC and male BBN-induced tumors. (B) Venn diagram showing the overlap of hypomethylated genes (top) and hypermethylated genes (bottom) between female human TCC and female BBN-induced tumors. (C) Pathway enrichment of hypomethylated and upregulated genes in male TCC using ENRICHR hallmark gene sets. Bar colors indicate -log10(p-values). (D) Pathway enrichment of hypermethylated and downregulated genes in female TCC. Bar colors indicate -log10(p-values).

RNA sequencing was performed on the same BBN tumor and normal tissue specimens used for DNA methylation analysis, comprising 8 male tumors, 2 male normal tissues, 6 female tumors, and 2 female normal tissues. Relative to normal bladder, male tumors exhibited 2,397 upregulated and 2,036 downregulated genes, whereas female tumors exhibited 1,645 upregulated and 1,381 downregulated genes (Supplementary Figures 6A and 6B). GSEA identified several programs in female BBN tumors that we also observed in human female tumors compared to their male counterparts, including EMT, mitotic spindle, and reduced UV response (Supplementary Figure 6C), supporting conservation of sex-associated transcriptional programs across species.

Integration of sex-stratified DMRs with DEGs identified one hypermethylated and downregulated gene, *Ace*, and 11 hypomethylated and upregulated genes in male BBN tumors (Supplementary Figure 7A). In female tumors, 17 genes were hypermethylated and downregulated, whereas *Sgk1* was the only hypomethylated and upregulated gene (Supplementary Figure 7B). Hypomethylated and upregulated genes in male tumors were enriched for UV response Up, TNF-α signaling, and glycolysis (Figure 6C), whereas hypermethylated and downregulated genes in female tumors were enriched for UV response Up, and xenobiotic metabolism pathways (Figure 6D).

We next assessed whether sex chromosome-associated features identified in human TCC were also evident in BBN-induced tumors. Application of the X-chromosome inactivation (XCI) escape gene signature revealed substantially higher XCI signature expression in female compared with male BBN tumors (Supplementary Figure 8A). Within the male tumors, application of the Y-chromosome gene signature identified two tumors with low Y-chromosome expression (BBN06 and BBN20), consistent with loss of Y-chromosome-associated gene expression (Supplementary Figure 8B).

To identify transcriptional differences associated with these sex chromosome signatures, we compared tumors with high and low XCI or Y-chromosome signature expression. Differential expression analysis identified *Apol8* as upregulated in the low-XCI group and *Sox5* as downregulated in the low-XCI group (Supplementary Figure 8C). In the Y-chromosome analysis, 7 genes were upregulated and 30 genes were downregulated in tumors with low compared with high Y-chromosome signature expression (Supplementary Figure 8D). Hallmark pathway analysis identified enrichment of adipogenesis and KRAS signaling among genes associated with the high Y-chromosome signature group (Supplementary Figure 8E). Together, these analyses demonstrated that sex chromosome-associated transcriptional features observed in human TCC could also be detected in BBN-induced tumors.

Together, these analyses demonstrate that BBN-induced bladder tumors recapitulate key tumor-associated DNA methylation and transcriptional features of human MIBC, including several sex-associated molecular programs. The greater conservation of male-associated methylation changes and preservation of selected sex-associated transcriptional programs support the BBN model for investigating epigenetic mechanisms underlying sex differences in bladder cancer.

### DNA methylation is associated with regulation of prognostic genes in bladder cancer

To assess the prognostic relevance of genes showing conserved methylation-associated expression across species, we analyzed the TCGA MSK 2020 bladder cancer cohort. Patients were stratified by upper and lower expression quartiles. High expression of *ONECUT2*, *PPP2R2C*, and *ADAMTS12* were significantly associated with shorter progression-free survival (Figure 7 A-C), supporting their prognostic relevance in MIBC.

**Figure 07.**
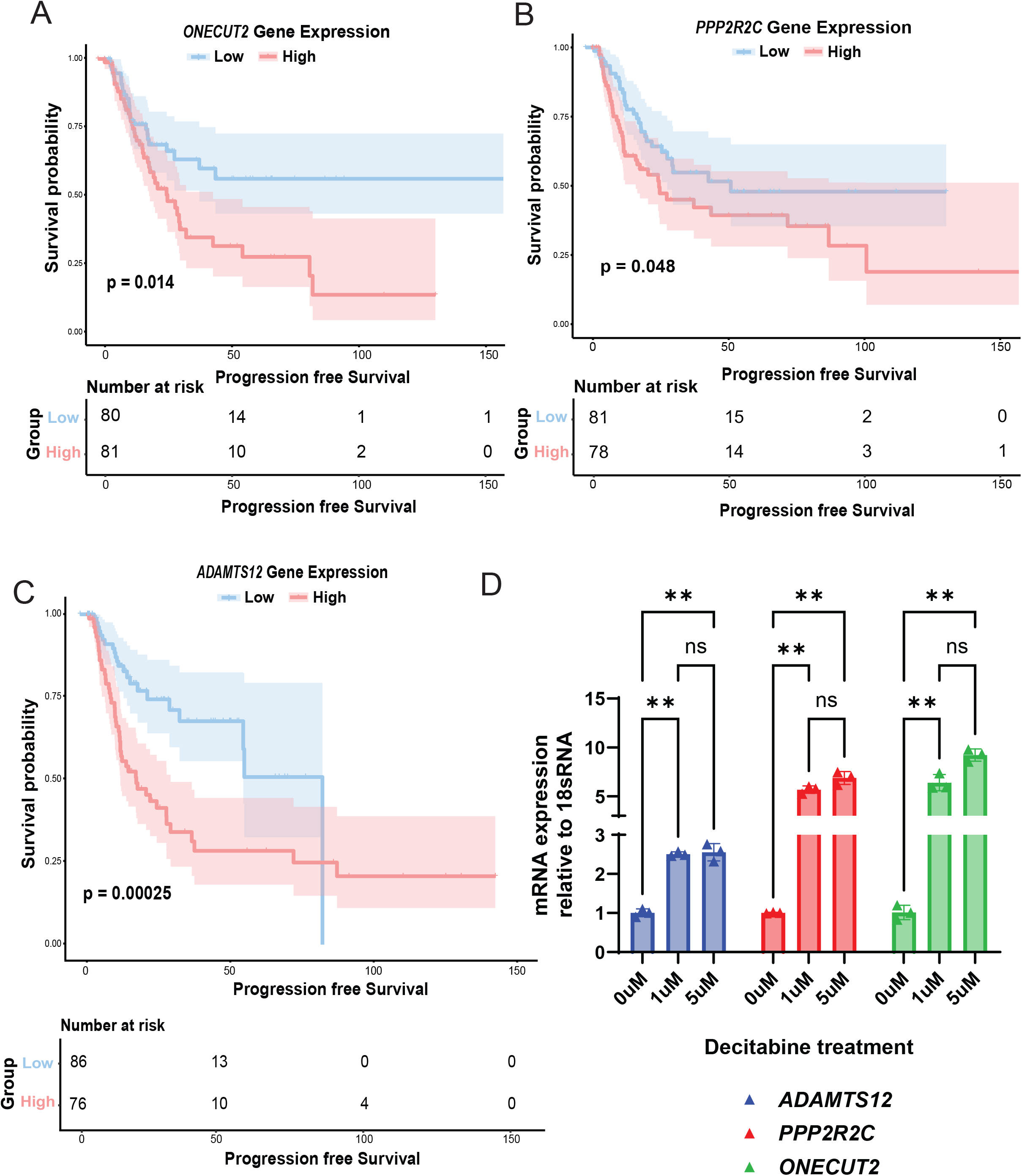
Genes regulated by DNA methylation play a prognostic role in bladder cancer. Progression-free survival for (A) ONECUT2, (B) PPP2R2C, and (C) ADAMTS12 was analyzed using the Survminer package in R. Patients were stratified into expression quartiles, with the 1st quartile as low and 4th quartile as high. Numbers at risk indicate patients remaining at each timepoint. Significant differences between low- and high-expression groups (p < 0.05) are shown. (D) qRT-PCR analysis showing expression changes of *ADAMTS12*, *PPP2R2C*, and *ONECUT2*, in the immortalized bladder cell line SV-HUC1 following decitabine treatment (0 µM, 1 µM, and 5 µM) for 48 hours.

To determine whether DNA methylation contributes to regulation of these genes, immortalized human bladder epithelial cells (SV-HUC1) were treated with the DNA demethylating agent decitabine for 48 hours. Decitabine significantly increased expression of *ADAMTS12*, *ONECUT2*, and *PPP2R2C* compared with vehicle treated controls (Figure 7D). Consistent with the activity of decitabine, DNMT1 protein levels decreased in a dose-dependent manner following treatment with 1µM and 5µM of decitabine (Supplementary Figure 9A). Together, these findings identify *ADAMTS12*, *ONECUT2*, and *PPP2R2C* as conserved, methylation-regulated genes whose expression is associated with shorter progression free-survival in MIBC.

## Discussion

Bladder cancer is more common in men, yet women experience worse outcomes^1,38^. Our sex-stratified analyses revealed marked epigenetic and transcriptional differences between sexes in MIBC. Male tumors displayed broader methylation remodeling and greater cross-species conservation, whereas female tumors, despite fewer DMRs, consistently showed enrichment of EMT and other tumor-associated programs. Consistent with our findings, a recent pan-cancer analysis of TCGA data identified BLCA as having a pronounced male-biased magnitude of cancer-associated DNA methylation changes^23^. The smaller number of female DMRs may reflect both the limited number of female samples and biological differences, as a similar disparity was observed in BBN tumors, where male and female cohort sizes were more comparable. The consistent enrichment of EMT-associated programs in female tumors across human and BBN datasets further suggests that female MIBC may exhibit distinct tumor-state biology despite less extensive methylation remodeling. Larger female cohorts with appropriate normal urothelial controls will be needed to distinguish these possibilities.

Our integrated methylation and transcriptomic analyses revealed widespread epigenetic remodeling in MIBC, with hypomethylated genes enriched in tumor-associated pathways including EMT and apical junction remodeling. Coordinated methylation and expression changes further support a role for DNA methylation in shaping tumor-associated transcriptional programs. Among conserved methylation-associated genes, *ADAMTS12*, *ONECUT2*, and *PPP2R2C* were hypomethylated and upregulated in both human and BBN tumors. ADAMTS12 contributes to extracellular matrix remodeling and metastasis^39–41^, while ONECUT2 acts as a transcriptional driver of aggressive disease states^42–44^. High expression of all three genes was associated with poorer progression-free survival, and pharmacologic DNA demethylation increased their expression, supporting methylation-dependent regulation. These findings identify conserved methylation-regulated genes with potential prognostic relevance in MIBC.

To determine whether findings in human MIBC were reflected in a relevant preclinical model, we evaluated the BBN mouse model focusing on the iUCSD subtype, which recapitulates the TCC phenotype represented in our human cohort^24,25^. BBN-induced tumors recapitulated over 7,000 DMRs observed in human MIBC and displayed conserved tumor-associated pathways and methylation-expression relationships. These findings extend the molecular similarity between human TCC and BBN-induced iUCSD tumors and support the use of this model for investigating epigenetic mechanisms in bladder cancer.

The sex-specific pathway patterns also provide potential insight into differences in tumor biology and response to cellular stress. Enrichment of the UV response pathway among methylation-associated genes likely reflects differences in cellular stress responses and DNA damage response programs rather than direct UV exposure. Notably, xenobiotic metabolism was enriched in female BBN tumors. This finding may be particularly relevant to the BBN model, as BBN carcinogenesis is influenced by metabolic activation, glucuronidation, and NRF2-regulated detoxification pathways^45^. The enrichment of xenobiotic metabolism in female BBN tumors may therefore reflect differences in carcinogen handling or preservation of a more differentiated urothelial state, although these possibilities require experimental validation.

Beyond autosomal differences, we developed an X-chromosome inactivation (XCI) escape gene signature to examine the contribution of sex chromosome gene expression to MIBC biology. Reduced expression of this signature was associated with poorer survival, after adjustment for sex and stage. Y-linked gene expression was not associated with survival, but low expression was associated with immune activation. Recent work has shown that Y chromosome loss in bladder cancer is accompanied by reduced expression of Y-linked genes and that this reduction becomes more pronounced with disease progression^46^. Y chromosome loss has also been associated with altered antitumor immunity and enhanced response to immune checkpoint blockade^47,48^. These findings implicate sex chromosome biology in distinct tumor states and clinical outcome in MIBC.

Sex chromosome-associated expression patterns were also evident in BBN tumors, including higher XCI escape signature expression in females and variation in Y-chromosome gene expression among males. Although these analyses were limited by the small number of tumors within individual signature groups, they suggest that the BBN model may provide a system for further investigating the contribution of sex chromosome regulation to bladder tumor biology.

Despite these strengths, several limitations warrant consideration. Cohort size, particularly for female TCC samples with matched NAs, was modest, which may limit detection of female-specific DMRs. In addition, male and female tumors were analyzed separately, and larger cohorts will be needed to directly compare the magnitude of these changes between sexes. Although BBN female tumors displayed similar patterns, confirmation requires larger, independent female cohorts. Functional validation focused on a subset of novel methylation-regulated genes, and the mechanisms underlying the regulation and function of *ADAMTS12*, *PPP2R2C*, and *ONECUT2* remain to be explored.

In summary, our study identifies sex-associated epigenetic and transcriptional programs in MIBC, methylation-regulated genes with prognostic significance, and substantial conservation of the human methylome in the BBN model. Male tumors displayed broader, conserved methylation remodeling, whereas female tumors showed distinct transcriptional programs including EMT. Together with the XCI escape gene signature, these findings deepen our understanding of sex-associated epigenetic regulation in MIBC and provide a framework for future biomarker and mechanistic studies.

## Materials and methods

### Cell culture

SV-HUC1 cells (ATCC, catalog no.CRL-9520) were cultured in F-12K media (Corning catalog no.: MSPP-F-12K) with 10% FBS (Corning catalog no.: 35-010-CV) and 1x Penicillin/Streptomycin (Corning catalog no.: 30-002-CI). Cells were maintained in 5% CO_2_ at 37^0^C. Cells were tested for Mycoplasma via MycoAlert Mycoplasma Detection Kit (Lonza, catalog no. LT07-218) after thawing. All experiments were performed with cells passaged less than 20 times after thawing.

### Decitabine treatment

SV-HUC1 cells (1 × 10⁶) were seeded in six-well plates. After cell attachment, cultures were treated with 0µM, 1µM, or 5µM of decitabine for 48 hours. A 100 µM stock solution of decitabine was prepared in sterile water and subsequently diluted in culture medium to achieve the indicated final concentrations.

### Western Blotting

Cells were lysed in Triton X-100/SDS lysis buffer (1% Triton X-100, 0.1% SDS, 50 mM Tris, 150 mM NaCl) containing protease (Sigma Aldrich catalog no.: 5892791001) and phosphatase inhibitors (Sigma Aldrich catalog no.: 4906845001). Protein concentration was determined using Bio-Rad protein assay (Biorad, Cat no. 500-0116). Equal amounts of protein lysates (50 µg) were resolved on a 4-20% gradient SDS-PAGE and electro transferred onto PVDF membranes. Blots were incubated with DNMT1 antibody (1:1000, Cell Signaling Technology Cat no. D63A6) and alpha tubulin antibody (1:1,000, Cell Signaling Technology Cat no. 1362) overnight at 4 °C, washed three times with TBST, incubated for 1 h at room temperature with LICOR IRDye anti-Rabbit (1:4000) antibody and LICOR IRDye anti-mouse (1:5000) antibody, washed three times with TBST, then detected and quantified on LICOR Odyssey Clx and ImageStudio software. Alpha tubulin was used for assessment of protein loading.

### RNA Isolation and cDNA Synthesis and qPCR

Total RNA was extracted using the Direct-zol RNA MiniPrep kit (Zymo Research, catalog no. R2052) and quantified using a Nanodrop 8000 system (Thermo Fisher Scientific). cDNA was synthesized from 500ng RNA using the iScript cDNA synthesis kit (Bio-Rad, catalog no.170-8891). SYBR green (BioRad, 172-5121) was used for qPCR with 10uM primers of *ADAMTS12*, *PPP2R2C*, and *ONECUT2* genes (Supplementary table 1).

### Carcinogen induced bladder cancer mouse model (BBN model)

Mice were housed in the Comparative Oncology Shared Resource (COSR) at Roswell Park, a limited-access barrier facility accredited by AAALAC International. 8 weeks old male and female C57BL/6 mice were administered with 0.1% BBN (TCI Chemicals, Cat # B0938) in their drinking water ad libitum until tumor collection. Bladder tumor formation was monitored through weekly abdominal palpations. High-grade disease was identified by the presence of palpable tumors and/or blood in the urine, at which point the mice were euthanized following Roswell Park Institutional Animal Care and Use Committee (IACUC) guidelines. Bladder tumors were removed and processed for histologic evaluation, RNA sequencing, and Illumina mouse methylation bead chip array.

### Histological evaluation of BBN tumors

BBN tumor tissues were fixed in 10% buffered formalin for 24 hours, processed for paraffin embedding, and sectioned at 5 microns on Starfrost adhesive slides (Mercedes Medical; Catalog #MER 7255/90/WH). Slides were deparaffinized in three xylene baths and rehydrated through graded alcohols (100% to 70%), followed by rinsing in distilled water. Hematoxylin and Eosin (H&E) staining was performed using the DAKO CoverStainer (Agilent Technologies, CS11830-2). After staining, slides were dehydrated through graded alcohol baths and cleared in three xylene baths before coverslipping. This process ensured high-quality staining for histological analysis by pathologist N. Sule.

### Roswell Park Comprehensive Cancer Center (Roswell Park) patient cohort

Tumor samples from patients with muscle-invasive bladder cancer (MIBC) were collected during radical cystectomy procedures at Roswell Park, with informed patient consent. De-identified tumors underwent RNA sequencing and Illumina 450k array for analysis. The tumors used in this study were collected and prepared according to the methods outlined in [Ramakrishnan et. al., 2017 and Ramakrishnan et. al., 2019] studies^18,27^.

### RNA sequencing and differential expression analysis

Paired-end RNA sequencing was performed on RNA isolated from BBN tumor samples (Male: Non-Tumor (NT) = 2, Tumor (T) = 9, Female: NT = 2, T = 8). RNA extraction was performed using a Qiagen miRNeasy mini kit (Cat#217044). RNA-seq analysis was conducted by Roswell Park Genomics Shared Resource. Sequencing libraries were prepared using the RNA HyperPrep Kit with RiboErase (Roche Sequencing Solutions) with 500 ng of total RNA, according to the manufacturer’s protocol. Initially, rRNA was depleted from the total RNA, followed by DNA digestion to remove gDNA contamination. The samples were then purified, fragmented, and primed for cDNA synthesis. Fragmented RNA was reverse transcribed into first-strand cDNA using random primers. The RNA template was then removed, and a replacement strand was synthesized, incorporating dUTP in place of dTTP to generate double-stranded cDNA. The ds cDNA was separated from the second strand reaction mix using Pure Beads (Kapa Biosystems), resulting in blunt-ended cDNA. An ‘A’ nucleotide was added to the 3’ ends of the blunt fragments, and multiple indexing adapters were ligated to the ds cDNA, preparing them for hybridization onto a flow cell. Libraries were amplified by PCR, purified, and validated for size using a 4200 TapeStation D1000 Screentape (Agilent Technologies, Inc.). The libraries were quantified via Kapa Biosystems qPCR kit, pooled at equimolar concentrations, denatured, and diluted to 350 pM, with 1% PhiX control added. The pool was loaded into a NovaSeq Reagent cartridge for 100-cycle paired-end sequencing on a NovaSeq6000, following the manufacturer’s guidelines (Illumina Inc.). Sequencing quality control was assessed using FastQC.

The raw reads that passed the quality filter from Illumina RTA were first preprocessed using FastQC^49^ for sequencing base quality control to be mapped to the human reference genome GrCh37 and the corresponding RefSeq annotation database using TopHat^50^. A second round of quality control using RSeQC^51^ was applied to mapped bam files to identify potential RNA-seq library preparation problems. From the mapping results, the read counts for genes were obtained by HTSeq^52^ using the intersection-strict option. Differentially expressed (DE) genes were identified using DESeq2 (v1.44.0)^53^. DESeq2 was performed on three groups for both human and BBN data: (A) Tumor vs Normals/NAs, (B) Male Tumor vs Normals/ NAs and Female Tumor vs Normals/NAs Differential, (C) Female vs male tumors. Expression rank order was utilized for subsequent Gene Set Enrichment Analysis (GSEA), performed using the clusterProfiler^54^ package in R. GSEA analysis was performed using the GSEA software, and the Hallmark pathway gene set was queried^55^.

### DNA methylation analysis in mouse and human bladder tissues

#### Mouse samples

DNA was extracted using the Qiagen QIAamp DNA Mini Kit (Cat# 51304) from 11 male (2 non-tumor [NT], 9 tumor [T]) and 10 female (2 NT, 8 T) mouse bladder samples. Genome-wide DNA methylation profiling was conducted using the Illumina Infinium Mouse Methylation BeadChip. A total of 500 ng of high-quality genomic DNA was bisulfite-converted using the Zymo EZ-96 DNA Methylation Kit (Zymo Research, Irvine, CA, USA), followed by amplification, fragmentation, purification, and hybridization onto the BeadChip per the manufacturer’s instructions. Arrays were washed, scanned using the Illumina iScan System, and processed at Roswell Park. Raw image data were analyzed using Illumina GenomeStudio V2011.1.

#### Data processing and DMR analysis

Minfi, a methylation array Bioconductor package, was used to read raw intensities, preprocess, and normalize values from methylation arrays^56^. Probes with detection P-values >0.01 were filtered out before preprocessing. preprocessQuantile was used to normalize raw data^56^. The M value was calculated using the formula M = log2((M+a)/(U+a)), where M represents the methylated intensity, U denotes the unmethylated intensity, and a is a constant offset (default value: a=100). Differential methylation analysis was performed separately on mouse and human methylation datasets. The R package DMRcate^57^ was utilized to identify and annotate differentially methylated regions (DMRs) between tumor and NAs tissues, as well as to assess sex-specific variations in DMRs. For the mouse data, we used the Illumina Infinium Methylation Array (mm10), and for the human data, the Illumina 450K array (hg19). Tumor versus normal/NAs tissue comparisons were conducted for each species using appropriate normalization and statistical testing pipelines (DMRcate using Limma), generating lists of DMRs defined by sets of adjacent probes showing significant methylation changes. DMRs were prioritized using Fisher’s multiple comparison statistics, incorporating both maximum differences (maxdiff) between CpG probes within a DMR and mean differences (meandiff), which calculate the average differences across CpG probes in each DMR. A Fisher statistic of P<0.05 was significant.

### Mapping of mouse DMRs to human genome

To map mouse DMRs to the human genome, the liftOver function from rtracklayer^58^ (v1.66.0) was run using the mm10ToHg19.over.chain file obtained from UCSC’s Lift Genomic Annotation database. Mouse DMRs were converted to putative orthologous human regions in hg19 coordinates. Only successfully mapped regions with unique mappings were retained for further analysis. Human and liftOver-transformed mouse DMRs were intersected using genomic coordinate overlap. Regions where mouse DMRs overlapped human DMRs were considered putatively conserved methylation loci between the two species. Genomic coordinates handling and overlap matching were performed using GRanges (v1.58.0)^59^.

For each overlapping DMR, we selected a single representative probe per region to visualize methylation patterns. The probe with the largest methylation value range (i.e., the largest difference between maximum and minimum β-values across all samples) within each DMR was chosen. This approach highlights the most informative sites per region. Methylation levels for these representative probes were then visualized in tumor vs. normal comparisons for both species to assess correspondence in methylation direction and magnitude.

### Multi-omics integration analyses

Maxdiff (from the methylation array data) and log2(Fold Change) (log2FC) (from the RNAseq expression data) statistics were utilized to quantify effect sizes. Gene symbols from the expression data were matched with their corresponding counterparts overlapping probe symbol(s) within a DMR. Since each DMR consists of many probes, the one with the largest probe-wise beta value range was picked as a representative for a given DMR. Sample β-values from that representative probe were extracted and matched with the corresponding sample DESeq2-normalized expression value. Spearman correlation was utilized to analyze the relationship between DNA methylation and RNA sequencing.

### XCI escape and Y chromosome gene signature

X-chromosome inactivation (XCI) escape and Y-chromosome were evaluated in human bladder tumor and BBN tumor samples. A set of 16 genes known to escape X-chromosome inactivation was identified through a literature search. The XCI gene signature included *KDM6A, ATRX, DDX3X, MAGEC3, EIF2S3, UTP14A, ALG13, CNKSR2, KDM5C, MID1, CA5B, GPM6B, JPX, SH3BGRL, SYP,* and *XIST*. Additionally, a set of 17 genes located in the male-specific region of the Y chromosome (MSY) and expressed in human bladder tissue was identified through a literature search^48,60^. The Y-chromosome gene signature included *DDX3Y, UTY, KDM5D, USP9Y, RBMY1A1, ZFY, TMSB4Y, EIF1AY, NLGN4Y, TBL1Y, HSFY2, PCDH11Y, PRY2, DAZ1, DAZ2, DAZ3,* and *DAZ*. Genes not present in the expression dataset were excluded from further analysis. Expression values were z-score normalized for each gene, and a composite signature score was calculated for each sample as the mean of scaled expression values across all genes in the signature using R. Samples were stratified into “High” and “Low” signature groups based on the median signature score. Gene signature scores were calculated using the singscore R package^61^, which employs a rank-based approach to quantify the expression of predefined gene sets in individual samples. Gene expression values were ranked using rankGenes^61^, and signature scores were computed using simpleScore^61^. The resulting signature scores and group classifications were appended to the phenotype metadata using the dplyr package for downstream analyses.

### Survival analyses

To evaluate the prognostic significance of the gene signature, survival analysis was conducted using clinical metadata. Overall survival status was derived by recoding patient vital status, where “DECEASED” was assigned as 1 (event) and “LIVING” as 0 (censored). Disease-free or overall survival time in months was used as the time variable. A Kaplan-Meier (KM) survival analysis was performed to compare survival between “High” and “Low” gene signature groups. The survfit function from the survival package was used to fit KM curves, and log-rank p-values were computed to assess statistical significance using the ggsurvplot function from the survminer package. For adjusted survival analysis, Cox proportional hazards models were fitted to assess the association between gene signature group and survival while controlling for sex and tumor stage (AJCC pathologic stage). Sex and AJCC stage were included as covariates in the multivariable model. AJCC stage was treated as an ordered factor when appropriate. Hazard ratios and confidence intervals were obtained using the coxph function of survival package in R. Adjusted survival curves were generated using survfit based on model predictions, holding covariates constant (e.g., sex = male), and visualized using ggsurvplot. Additionally, interaction terms between signature group and sex were included to assess joint effects on survival. Survival outcomes were stratified by combined sex and signature group levels to visualize differential survival patterns across subgroups.

### Statistical Analysis

Quantitative data were compared among three groups using one-way analysis of variance (ANOVA). A P value of less than 0.05 was considered statistically significant.

## Study approval

All human tissue samples and associated clinical data used in this study were obtained under protocols approved by the Institutional Review Board of Roswell Park Comprehensive Cancer Center. Written informed consent was obtained from all participants prior to participation. All animal studies were conducted in accordance with protocols approved by the Institutional Animal Care and Use Committee of Roswell Park Comprehensive Cancer Center and in compliance with institutional and applicable regulatory guidelines.

## Data availability

The RNA-sequencing and DNA methylation array datasets generated and analyzed in this study will be deposited in the Gene Expression Omnibus (GEO) and made publicly available upon publication. The supporting bioinformatics and statistical analysis code used in this study will be made available from the corresponding author upon request.

## Acknowledgements

We thank Spencer Rosario for providing critical insights and feedback during manuscript preparation. We also thank Sarah Athans for critical reading and providing comments on the manuscript. We acknowledge the DOD W81XWH-19-1-0636 (Foster); Roswell Park Alliance Foundation (Foster and Huss); National Cancer Institute (NCI) grant P30CA016056 involving the use of Roswell Park Comprehensive Cancer Center’s Shared Resources.

## Author Contributions

All authors read and approved of the final manuscript. Developed analysis tools and analyzed data: BD, ECG and AW. Writing manuscript: BD and AW. ECG, BF, JW, and WH contributed to the development and writing methods.

## Notes

### Competing Interest Statement

The authors have declared no competing interest.

