## Supplementary figures for "Sex Differences in DNA Methylation and Gene Expression Reveal Distinct Molecular Programs in Muscle-Invasive Bladder Cancer"

### Supplementary figure panels

### Supplementary Figure 01

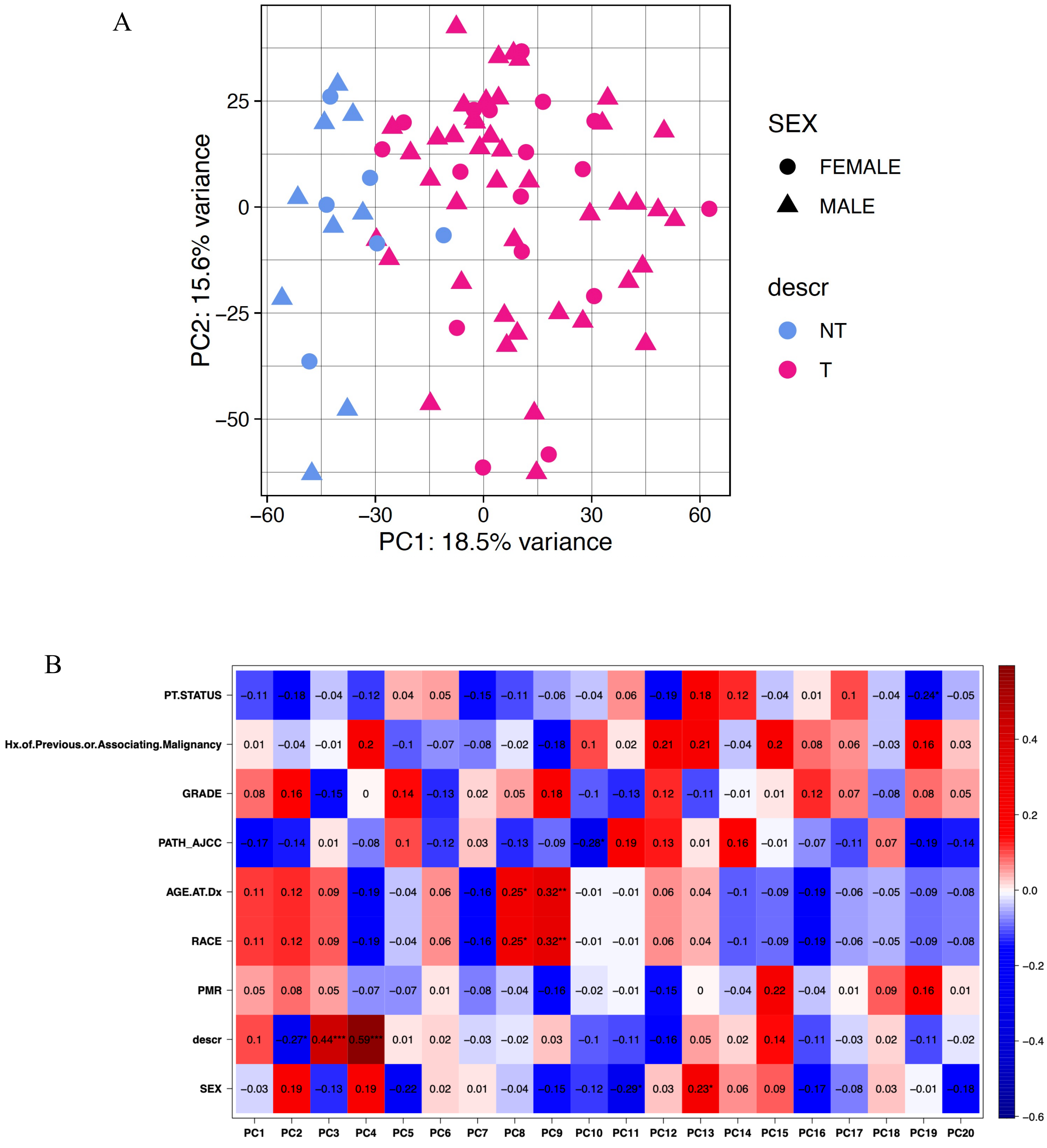

### Supplementary Figure 02

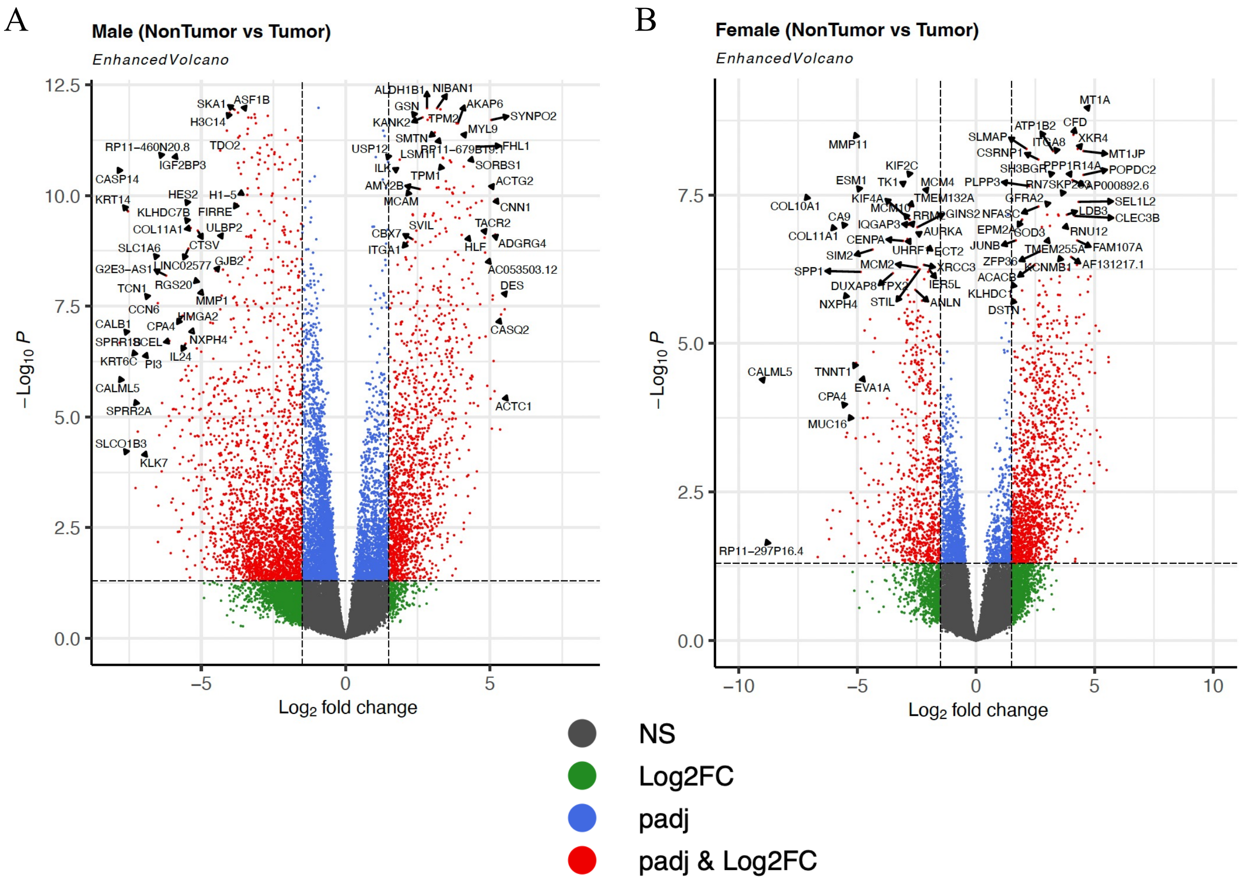


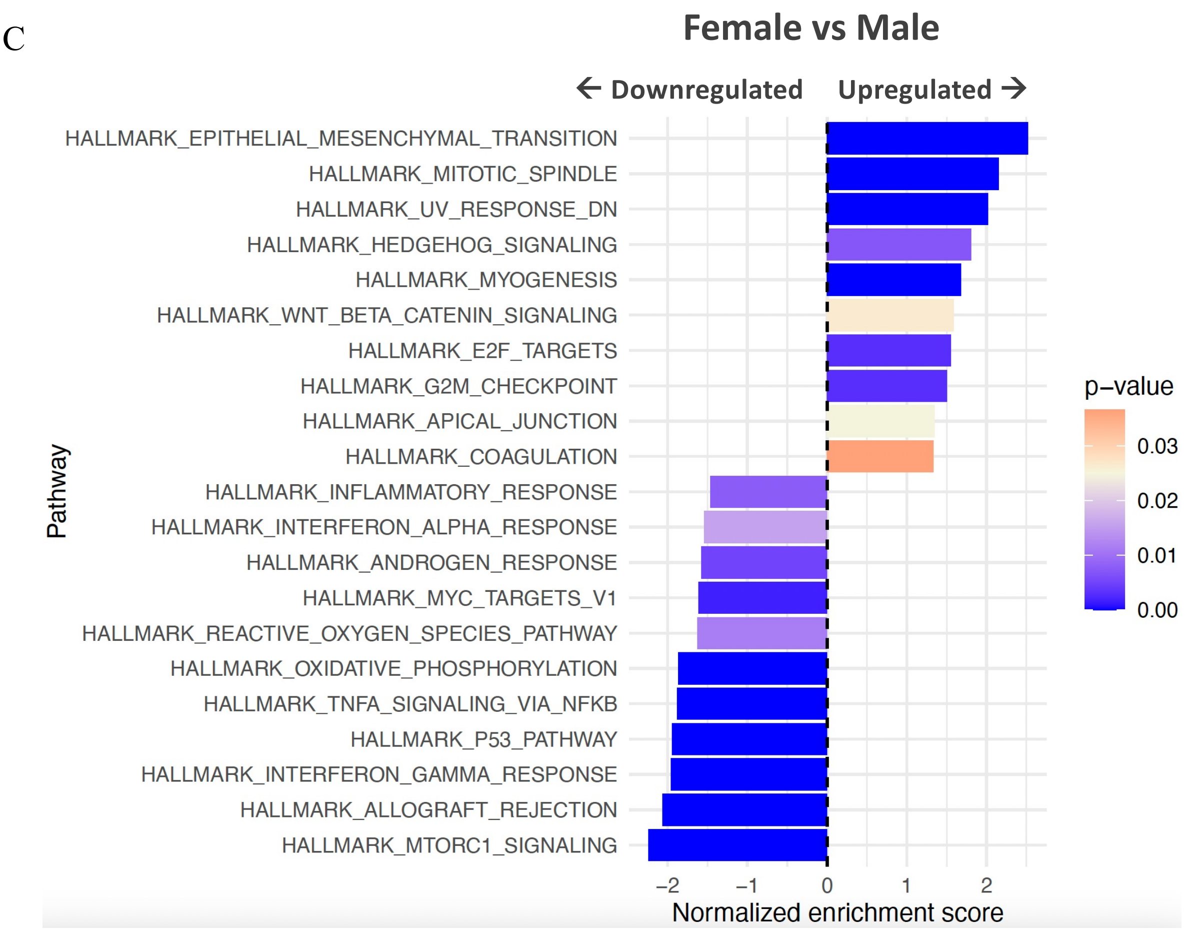

### Supplementary Figure 03

A

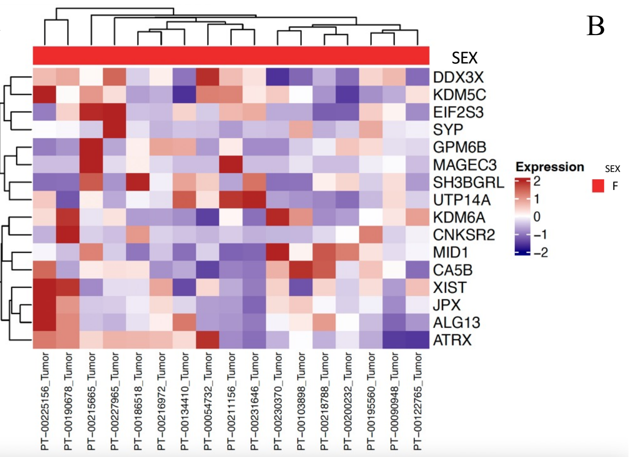


B

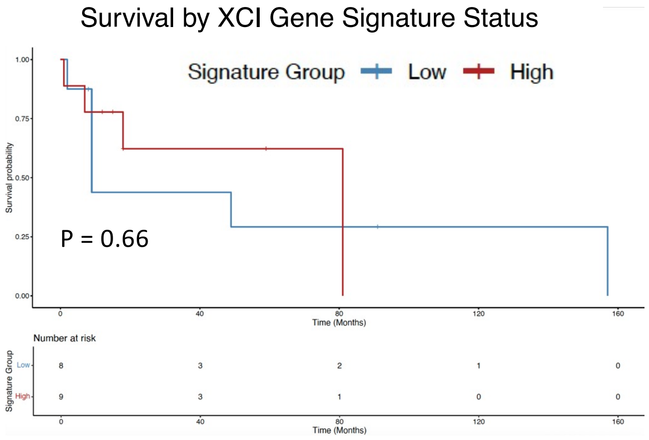


C

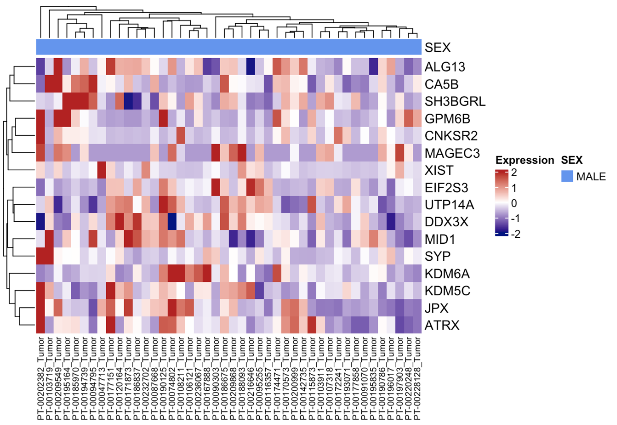


D

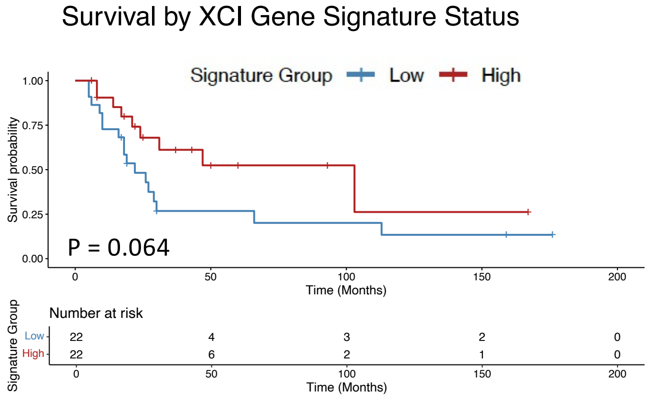

### Supplementary Figure 04

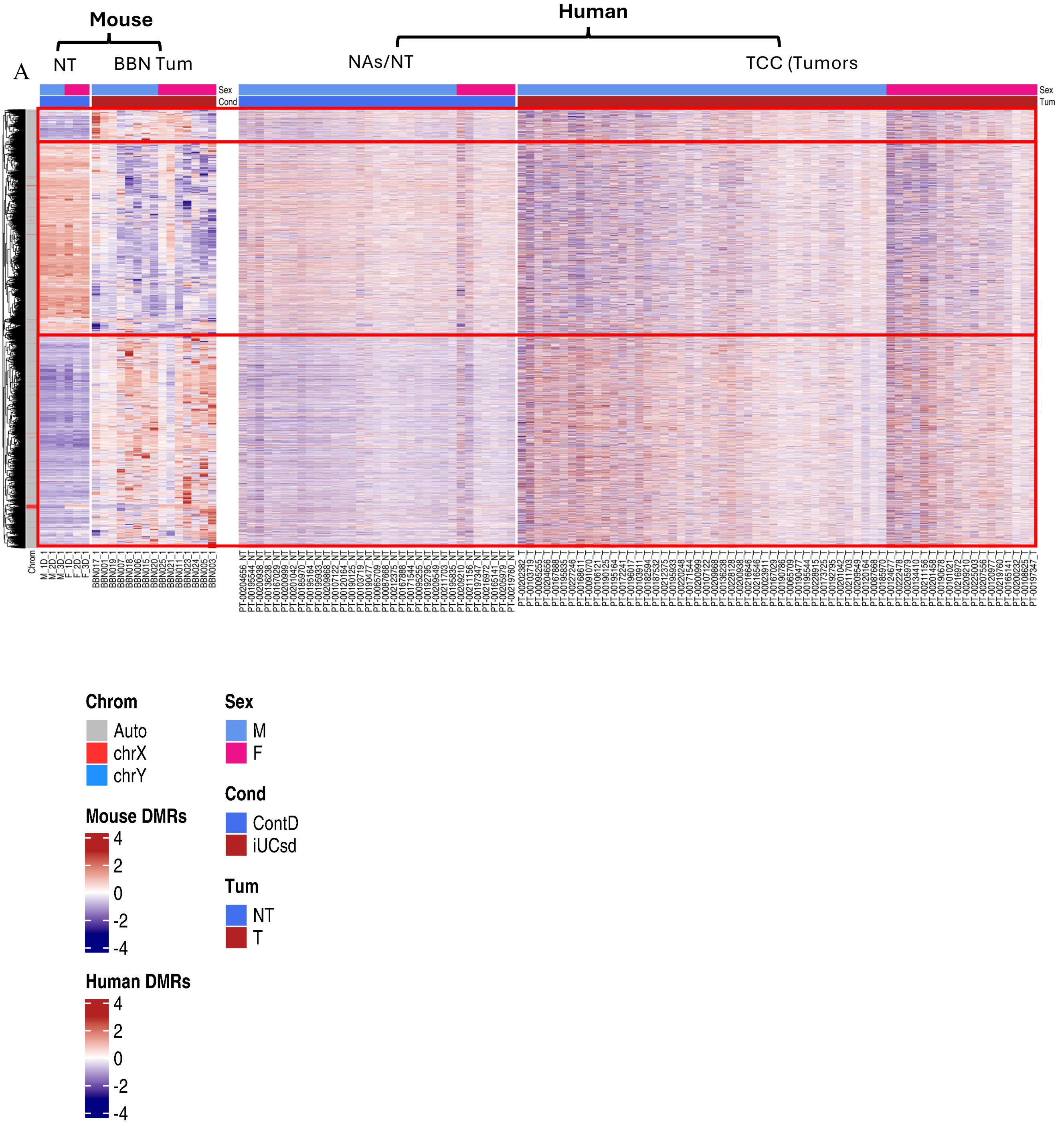

### Supplementary Figure 05

A

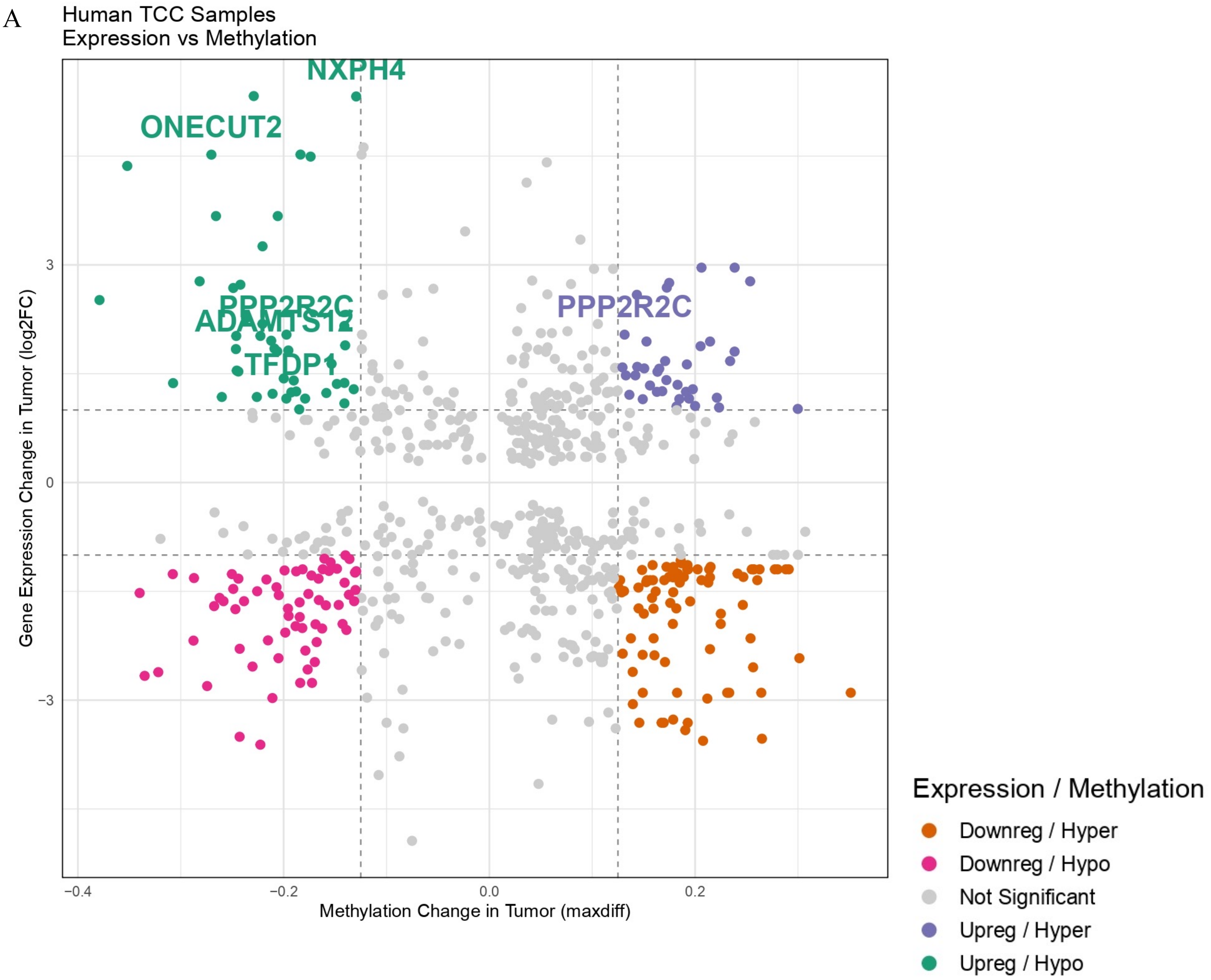


B

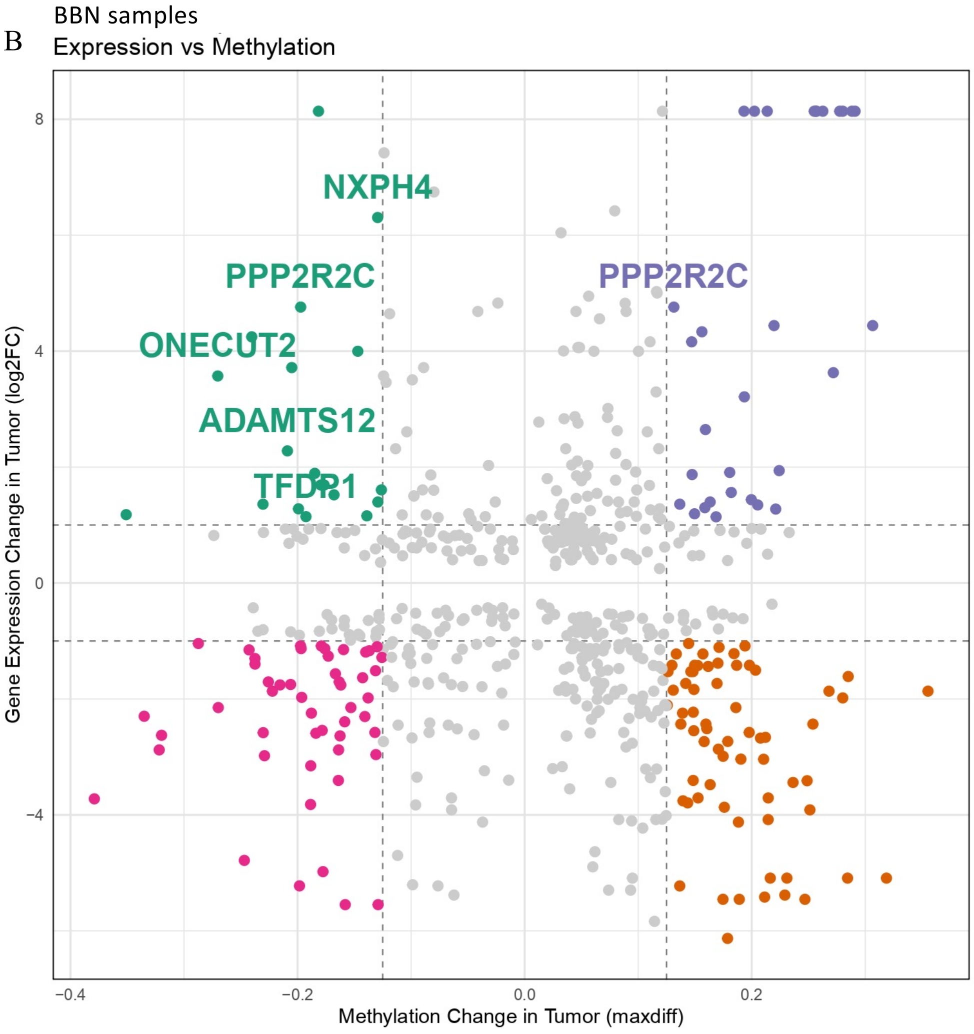

### Supplementary Figure 06

A

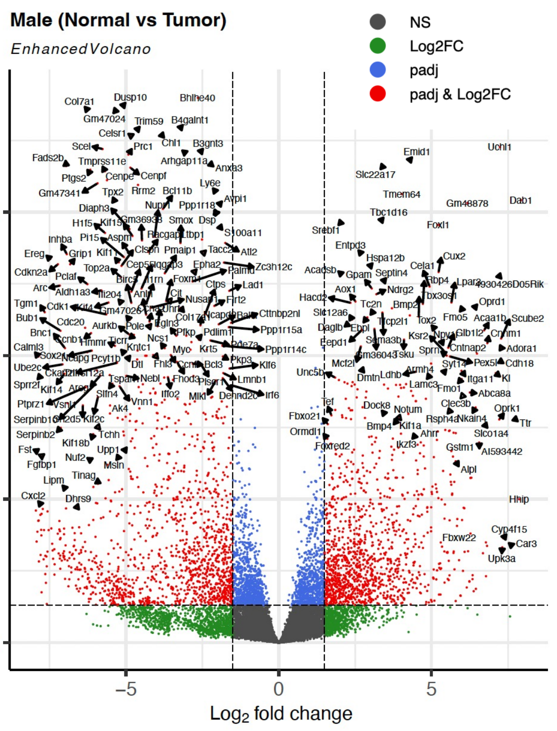


B

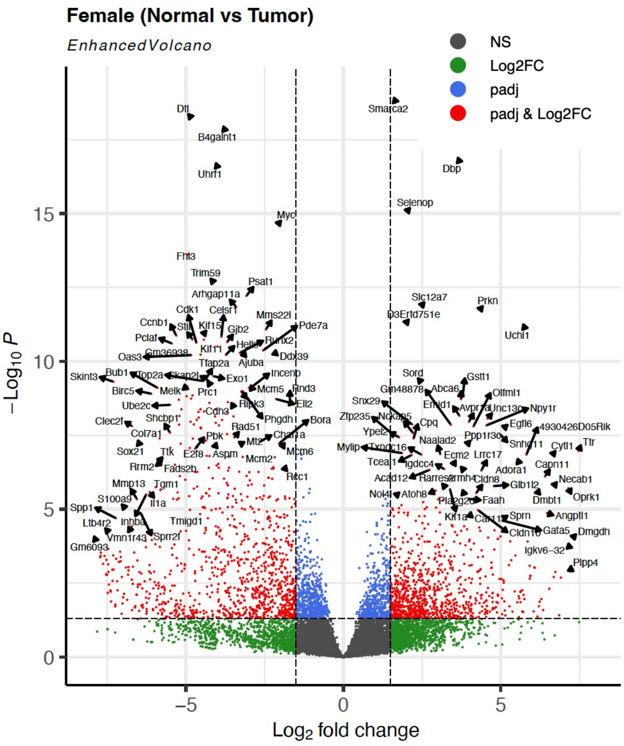


C

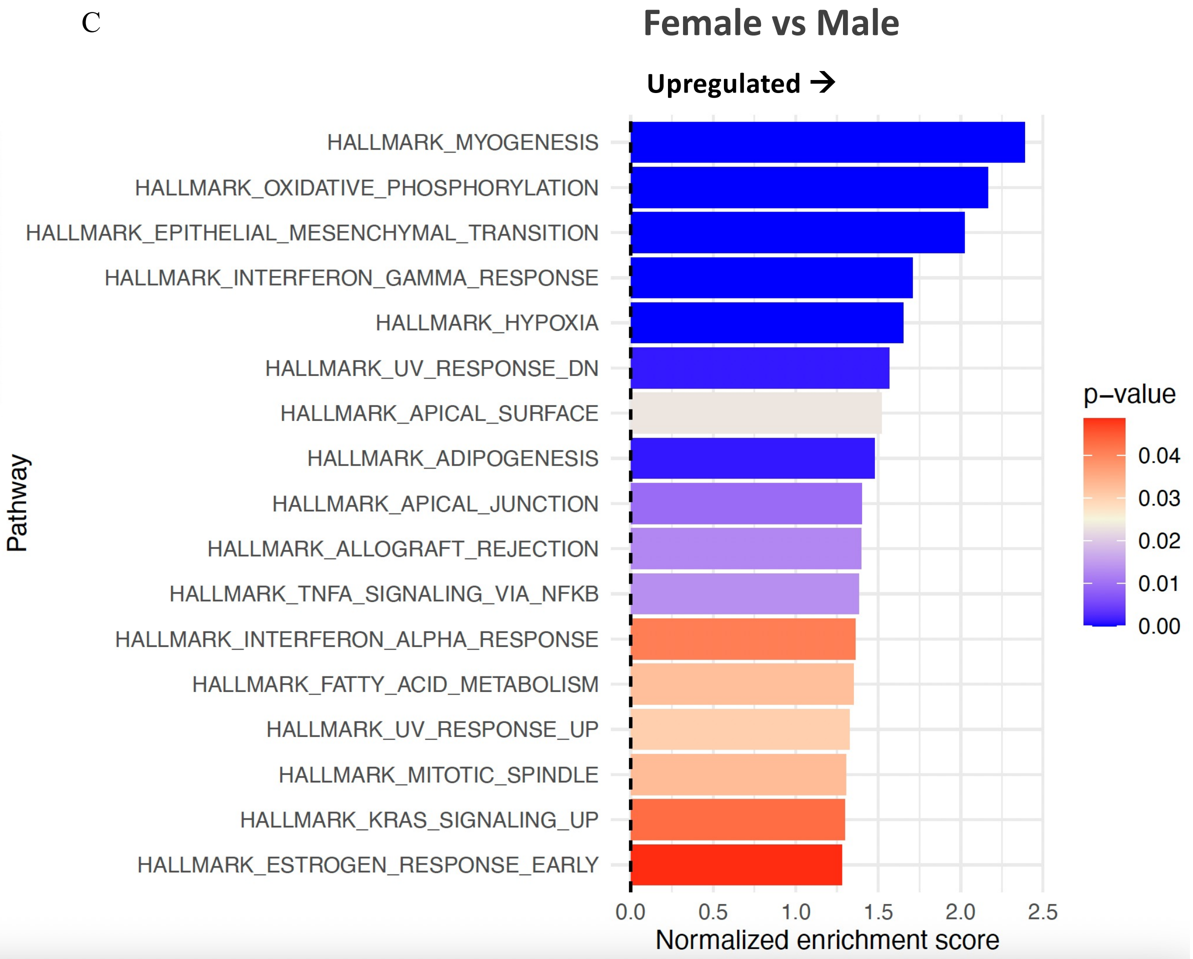

### Supplementary Figure 07

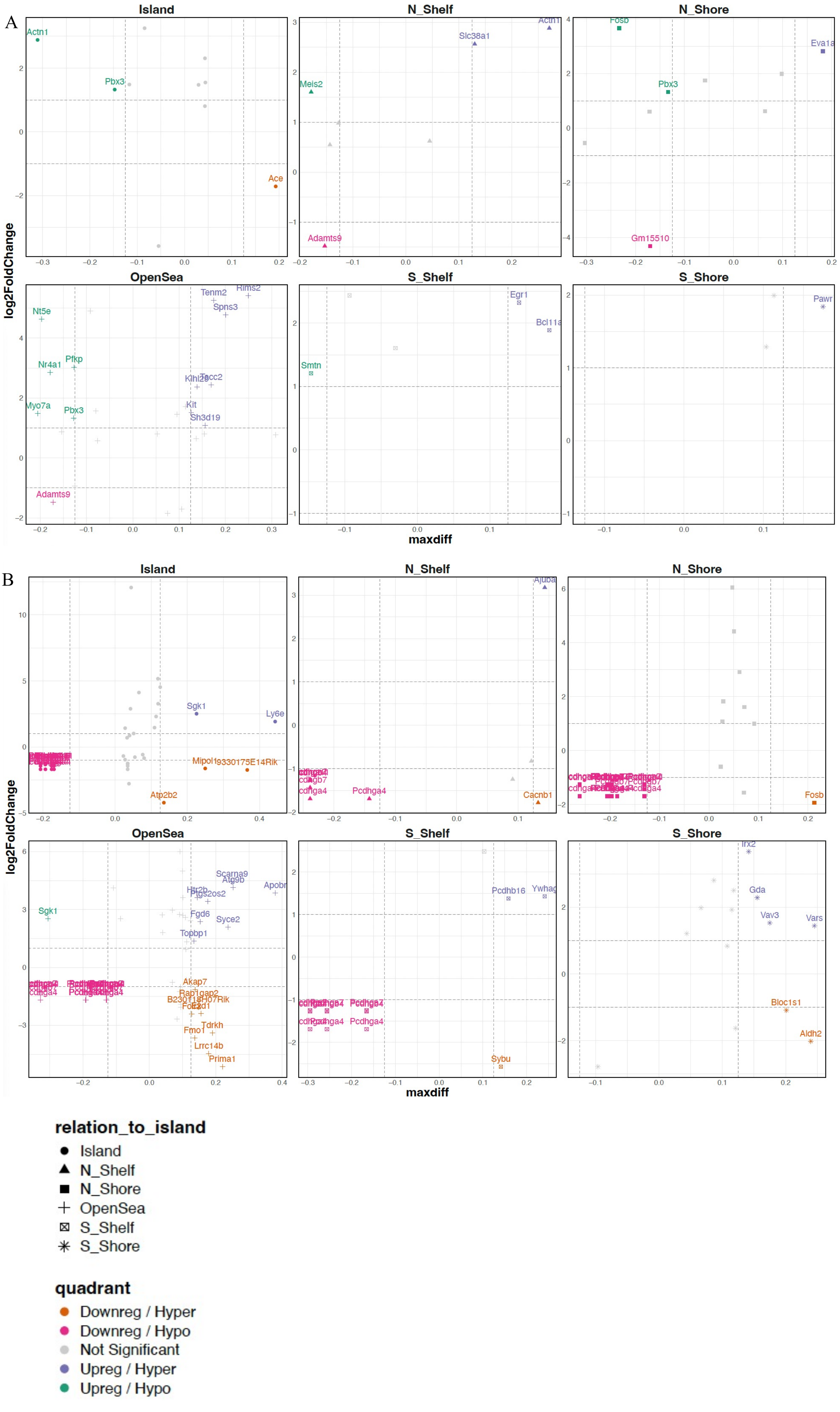

### Supplementary Figure 08

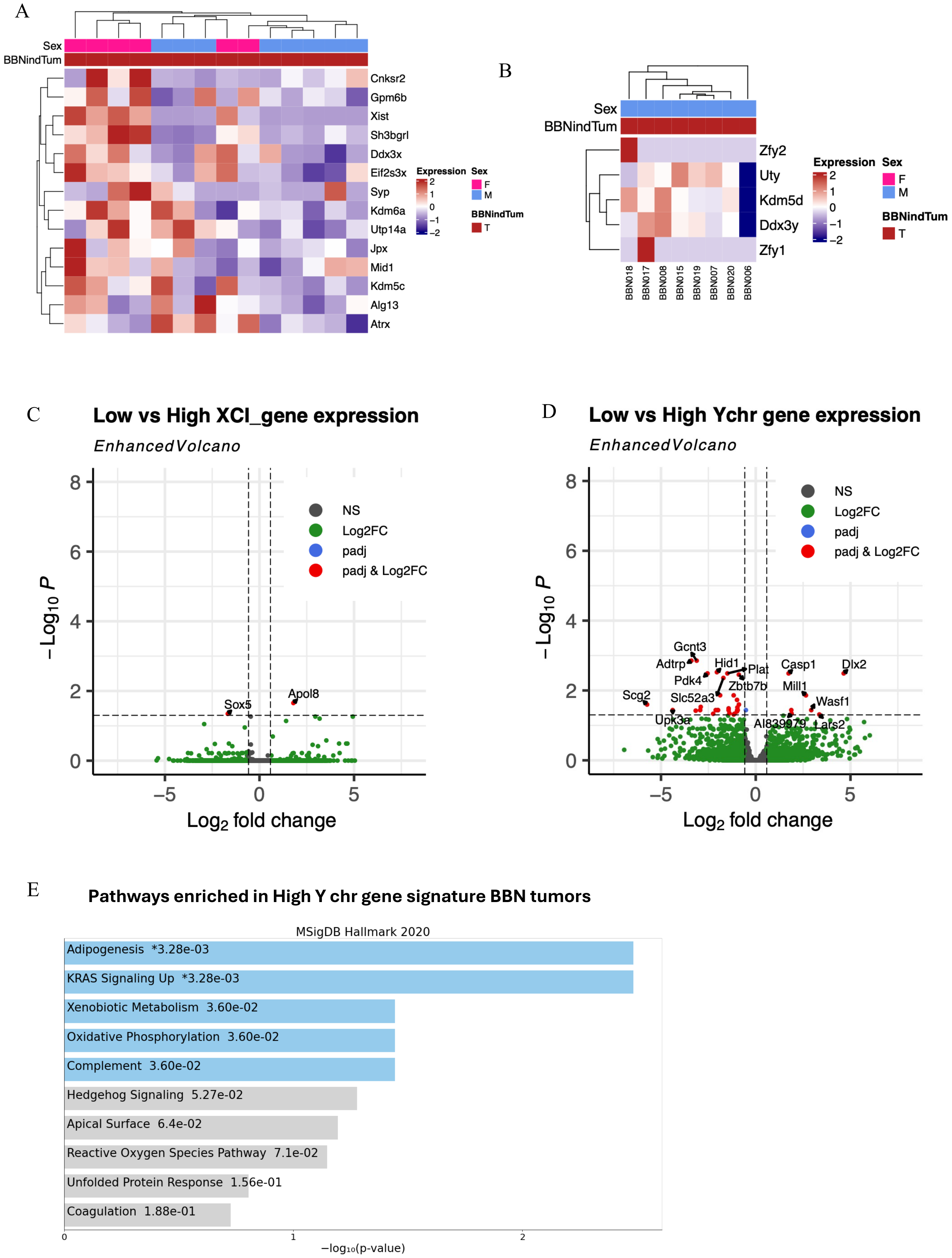

### Supplementary Figure 09

A

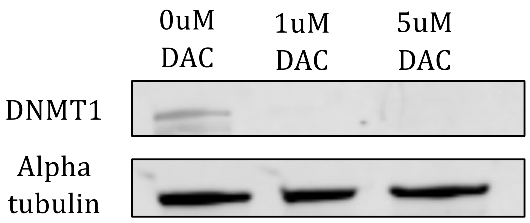
